# Integrative Structure of Norovirus NS3 Suggests a Role in RNA Transport

**DOI:** 10.1101/2025.06.17.659504

**Authors:** Meryl Haas, Jake T. Mills, Cynthia Kelley, Arnab Das, Henrietta Hóf, Coco Niemel, Tim Donselaar, Ieva Drulyte, Frank J.M. van Kuppeveld, David Dulin, Joost Snijder, Morgan R. Herod, Daniel L. Hurdiss

## Abstract

Human noroviruses (HuNoVs) are the leading global cause of non-bacterial gastroenteritis, yet no vaccines or antiviral therapies are currently approved^1^. The viral non-structural protein NS3 is a membrane-bound AAA+ ATPase of superfamily 3 (SF3) with multiple proposed roles in the norovirus replication cycle. However, the structure of NS3, and the mechanisms by which it contributes to genome replication and membrane remodeling, have remained unknown. We engineered a soluble, hexameric, and catalytically active form of NS3 and determined its cryo-EM structure in the presence of a nucleotide analogue at 2.9 Å resolution. The structure adopts a split lock-washer architecture characteristic of AAA+ motors that operate via a hand-over-hand translocation mechanism^2^. Modeling of the nucleotide-binding site reveals conserved features governing ATP binding and hydrolysis. Using integrative modeling with AlphaFold3, we generated a full-length, membrane-associated model of NS3, in which an N-terminal transmembrane domain, central helical bundle, and AAA+ motor form a continuous conduit. This model supports a role for NS3 as a candidate membrane-spanning RNA translocase that couples ATP hydrolysis to genome movement. This structural framework helps address long-standing gaps in our understanding of norovirus replication and establishes a basis for mechanistic studies and structure-guided antiviral design.

## INTRODUCTION

HuNoVs pose a substantial health burden to society, contributing to ∼700 million infections and over 200,000 annual deaths. These viruses are the primary cause of non-bacterial gastroenteritis, often referred to as “stomach flu” or “stomach bug”, marked by symptoms such as sudden onset diarrhea and vomiting^1,3,4^. Compounding this impact, the global economy suffers an approximate cost of €60 billion annually, encompassing expenses related to lost workdays and healthcare expenditures linked to these infections^5^. Transmission of the virus occurs primarily through the fecal-oral route, facilitated by contact with infected individuals or exposure to contaminated food, water, or infectious aerosols generated by vomiting^4^. This high infectivity and efficient transmission give rise to the potential for newly emerging norovirus strains to spark worldwide epidemics^6^. Notably, despite these risks, there are currently no approved vaccines or antiviral treatments targeting noroviruses^1,7^.

Noroviruses are non-enveloped, positive-sense single-stranded (ss) RNA viruses classified within the genus *Norovirus* and the family *Caliciviridae*. Together with enteroviruses, another notable group of human pathogens, noroviruses belong to the order *Picornavirales*, a diverse assemblage of RNA viruses that infect animals, protists, and plants. Within the genomes of these viruses, there is a characteristic three-domain replication block (Hel-Pro-Pol), which comprises a putative <u>hel</u>icase belonging to the SF3 clade of AAA+ ATPases, a chymotrypsin-like <u>pro</u>teinase and a superfamily I RNA-dependent RNA <u>pol</u>ymerase^8,9^. The norovirus protease and polymerase, NS6 and NS7, have been studied by X-ray crystallography and the molecular basis for their enzymatic activity is understood^10,11^. However, no such information is available for the norovirus NS3, which has been implicated as having multiple functions within the norovirus replication cycle, including: i) ATP-dependent helicase activity^12^, ii) ATP-independent RNA chaperoning^13^, iii) rearrangement of intracellular membrane alterations during replication organelle formation^14,15^, and iv) the induction of apoptosis^16^.

SF3 helicases typically assemble into hexameric rings and couple ATP hydrolysis to conformational changes that generate mechanical force on nucleic acid substrates^2^. These enzymes possess three conserved motifs essential for nucleotide binding and hydrolysis, Walker A, Walker B, and motif C, all of which are present in norovirus NS3^17,18^. In DNA viruses, SF3 proteins function as replicative helicases, and several oligomeric structures have been resolved^2,19–23^. However, no functional oligomeric assembly of an SF3 protein from an RNA virus has been structurally characterized to date, although crystal structures of picornaviral 2C AAA+ domains have been reported^24–26^. This has left a gap in our mechanistic understanding of the replication machinery employed by positive-strand RNA viruses.

Here, we present the first structure of the norovirus NS3 protein in a nucleotide analog-bound state. Through integrative modeling, we propose that NS3 functions as a transmembrane RNA translocase, coupling membrane-associated genome replication to RNA transport. This insight may fundamentally reshape our understanding of SF3 protein function in positive-strand RNA viruses. Our findings not only open new avenues for exploring how noroviruses replicate and package their genomes, but also provide a structural blueprint for targeting NS3 in future antiviral strategies.

## RESULTS

### Engineering a soluble and enzymatically active NS3 hexamer

Sequence homology with known SF3 proteins and predictive modeling indicates that NS3 comprises an N-terminal membrane-binding domain (NTD), a helical bundle domain (HBD), and a C-terminal AAA+ ATPase domain containing the signature Walker A, Walker B, and motif C required for nucleotide binding and hydrolysis^9^. Previous studies on homologous picornavirus 2C proteins have shown that their N-terminal amphipathic helix renders recombinant proteins poorly soluble but is necessary for efficient formation of enzymatically active hexamers; this requirement has thus far precluded structural studies of SF3 protein quaternary structures from RNA viruses^27,28^. Previously, we demonstrated that enzymatic ATPase activity can be restored in picornaviral 2C protein when arranged in a hexameric complex by replacing the NTD with a synthetic hexamerization domain^26^.

To rationally engineer a soluble recombinant HuNoV NS3 construct that retains the ability to form functional oligomers, we first generated AlphaFold2 predictions of the hexameric complex for HuNoV. These models indicated a suitable truncation site between the NTD and HBD, resulting in a construct lacking the first 61 amino acids of NS3, annotated as ΔN61 (Supplementary Figure 1A). We then inserted a soluble hexamerization domain, cc-hex^29^, and a linker region before our ΔN61 NS3 construct. To facilitate expression and purification, we added an N-terminal hexa-histidine tag (His6) and an MBP-tag to the N-terminus of our hex-ΔN61-NS3 construct, separated by a 3C protease cleavage site (MBP-hex-NS3, Figure 1A, Supplementary Figure 1B). For comparison, we also generated the equivalent ΔN61 NS3 construct without the hexamerization domain (MBP-NS3).

**Figure 1:**
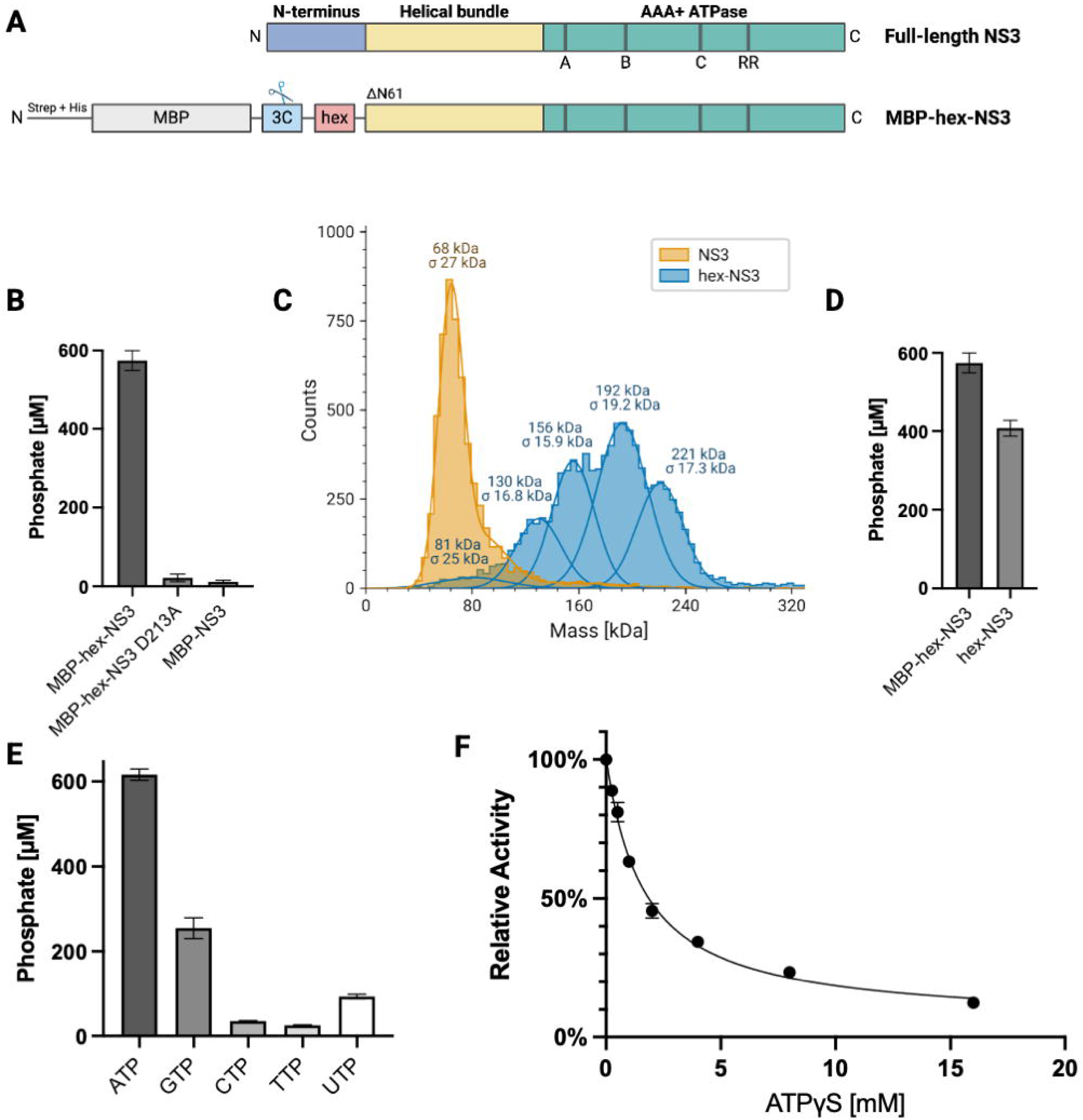
Engineering a soluble, enzymatically active HuNoV NS3 hexamer. (A) Linear representation of the full-length NS3 protein (top) and the engineered hex-NS3 construct (bottom). The full-length NS3 protein comprises a membrane-binding N-terminal domain (NTD), a helical bundle domain (HBD), and an AAA+ ATPase domain comprising Walker A, B, motif C, and arginine fingers (RR). In hex-NS3, the NTD (residues 1-61), is replaced by the 32-residue cc-hex coiled-coil sequence. (B) ATPase activity for the WT MBP-hex-NS3, the D213A mutant, and WT MBP-Δ61-NS3. (C) Mass photometry analysis of Δ61-NS3 and hex-Δ61-NS3. (D) ATPase activity for the MBP-hex-NS3 and following removal of the MBP. (E) NTPase activity of hex-NS3 with different nucleotides. (F) ATPase activity of hex-NS3 in the presence of increasing concentrations of ATPγS. For panels B, D, E, and F, representative data from one of two independent experiments, each performed in technical duplicate, are shown. Error bars represent the standard deviation of the two technical replicates.

Following expression in E. coli and purification (Supplementary Figure 2), we tested the ATPase activity of our MBP-tagged NS3 constructs using the malachite green assay, which detects free phosphate produced by nucleoside triphosphate hydrolysis. To confirm NS3-specific ATP hydrolysis, we substituted the critical Walker B residue D213 with alanine, which abolished ATPase activity (Figure 1B). Similarly, no detectable enzymatic activity was observed for the monomeric construct, highlighting the importance of oligomerization for nucleotide triphosphatase activity.

Subsequent removal of the MBP-tag demonstrated that both the monomeric and hexameric constructs remained soluble. To assess the oligomeric states of our NS3 constructs, we employed mass photometry and native mass spectrometry analyses. While NS3 in the absence of the cc-hex domain (33.6 kDa) was found to form primarily monomers and dimers, hex-NS3 (38.2 kDa) formed higher oligomers ranging from trimers to hexamers, highlighting the importance of the N-terminal region for oligomerization in NS3 (Figure 1C and Supplementary Figure 3). We then confirmed that the untagged hex-NS3 retains ATPase activity (Figure 1D). Due to its slightly higher activity and less involved purification protocol, the MBP-tagged construct was used for future NTPase assays described in this study.

The nucleotide specificity of NS3 was tested by providing different nucleoside triphosphates (ATP, GTP, CTP, TTP, and UTP) as substrates. NS3 exhibited a preference for ATP, followed by GTP, with ∼3-fold lower activity (Figure 1E). Notably, this contrasts with earlier studies that reported minimal discrimination between nucleotides by NS3^12^. As a prelude to structural analysis, we assessed whether Adenosine 5′-O-(3-Thiotriphosphate) (ATPγS) could compete for binding to the orthosteric site and thus serve as a suitable nucleotide analogue to capture the *holo* complex by cryogenic Electron Microscopy (cryo-EM). Our results demonstrated a dose-dependent decrease in ATP hydrolysis in the presence of increasing concentrations of ATPγS (Figure 1F), underscoring its utility for structural studies.

### Cryo-EM analysis of the NS3 hexamer

To assess the suitability of our hex-NS3 construct for structural studies, we first evaluated sample quality using negative stain electron microscopy (EM). Prior to grid preparation, hex-NS3 was incubated with ATPγS to stabilize the nucleotide-bound state. Micrographs revealed monodisperse, ring-shaped particles consistent with a hexameric assembly. In addition, we infrequently observed filamentous structures (Supplementary Figure 4A). Subsequent 2D classification of the ATPγS-bound sample yielded class averages consistent with the expected oligomeric architecture (Supplementary Figure 4B).

We next prepared cryo-EM grids and determined a three-dimensional reconstruction of the ATPγS-bound hex-NS3 complex at a global resolution of 2.9 Å (Figure 2A and Supplementary Figures 5-6). The resulting structure shows that six copies of NS3 assemble into a funnel-shaped hexamer approximately 80 Å in height. The HBD and AAA+ ATPase domain form upper and lower tiers measuring ∼70 Å and ∼105 Å in diameter, respectively. Unlike the C6-symmetric model predicted by AlphaFold2 (Supplementary Figure 1A), our cryo-EM reconstruction adopts a split lock-washer conformation, previously described in other AAA+ ATPases^2^. The six subunits form a right-handed spiral with a rise of ∼1.5 Å per protomer and a gap of ∼6.5 Å between the topmost and bottommost subunits. Notably, the HBDs make minimal, if any, lateral contacts with each other.

**Figure 2:**
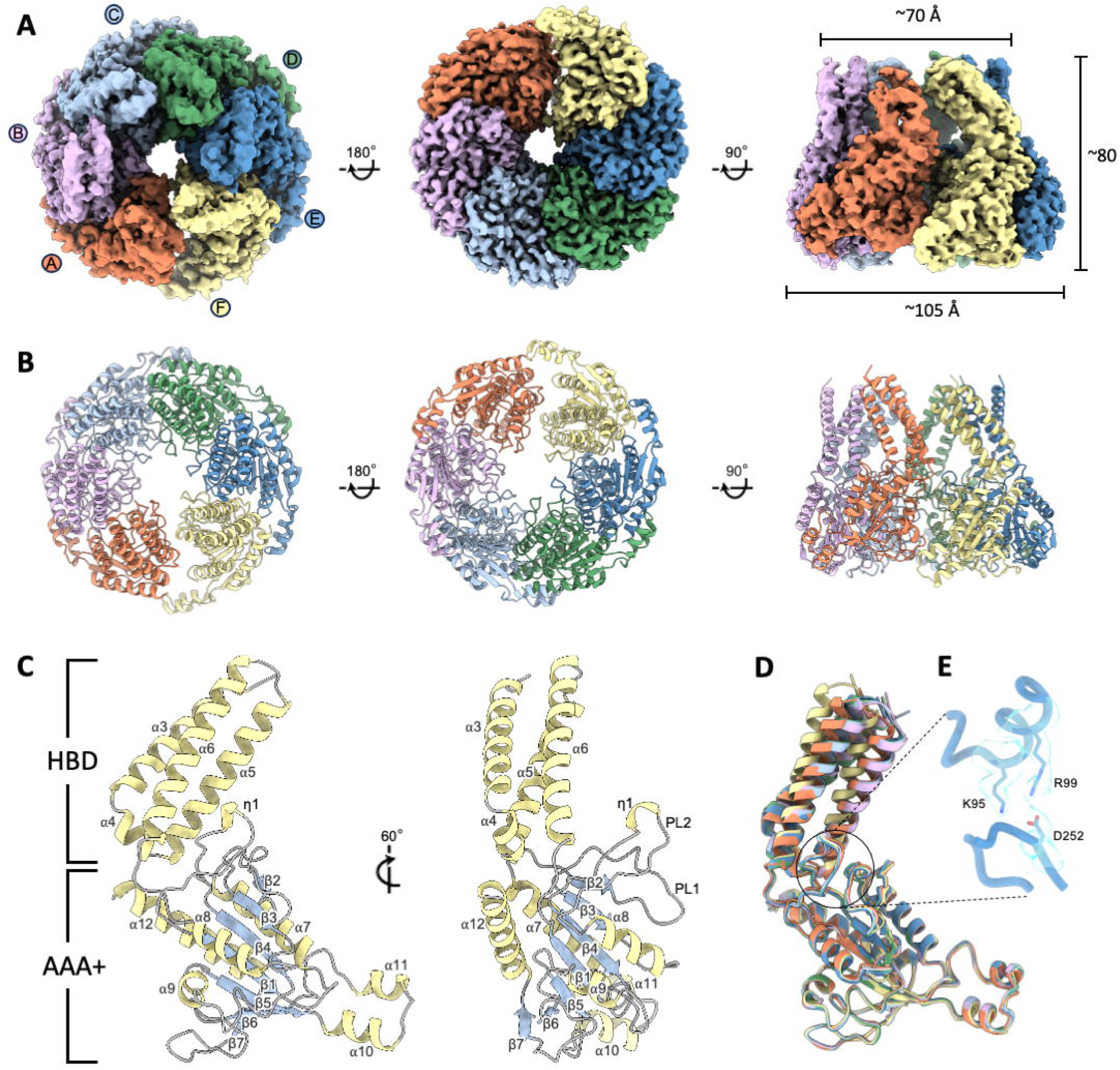
The architecture of the ATPγS-bound NS3 hexamer in a split lock-washer conformation. (A) Three orthogonal views of the EM map of the NS3 hexamer with individual protomers shown in different colors. (B) As shown in (A) for the atomic model. The ∼6.5 Å gap between protomers A and F is apparent in the first top view. (C) Cartoon representation of subunit C with helices in yellow, strands in blue, and loops in gray. The HBD and AAA+ domain have been labeled. (D) Superposition of the six NS3 protomers. (E) Zoomed-in view of the interface between the HBD and AAA+ domains in subunit E, with residues involved in salt bridge formation shown as sticks and the corresponding EM density shown in blue.

Next, we generated an atomic model encompassing residues 64–363 across all six subunits (Figure 2B and Supplementary Figure 7), as well as 204 water molecules. At the tertiary structure level, our cryo-EM model agrees closely with AlphaFold2 predictions. The RMSD across all aligned atoms ranged from 3.2 to 5.4 Å, with even better agreement for the AAA+ domain alone (RMSD 0.9–1.2 Å). Structurally, the HBD adopts a four-helix bundle (residues 64–148), connected to the AAA+ domain by a five-residue linker (Figure 2C). As expected, residues 154–363 form a canonical Rossmann fold, with all conserved SF3 motifs well-resolved. We also modelled two pore loops: residues 192–207 (pore loop 1) and residues 230–252 (pore loop 2).

Structural homology analysis using Foldseek and DALI confirmed strong similarity between the AAA+ domain and picornaviral 2C proteins, papillomaviral E1 helicases, and Rep helicases from single-stranded DNA viruses. For the NS3 HBD, AtBAG1, a BAG-family co-chaperone from *Arabidopsis thaliana*, was identified as the closest experimentally determined structural homolog^30^. To further explore the evolutionary relationship between NS3 and other viral SF3 helicases, we conducted structure-based phylogenetic analysis using available high-resolution structures of AAA+ domains from SF3 helicases. This analysis showed that NS3 clusters closely with 2C proteins from members of the family *Picornaviridae* (Supplementary Figures 8), reinforcing their shared ancestry and structural conservation within the order *Picornavirales*.

Finally, RMSD calculations across all six protomers revealed only minor angular deviations in the HBD–AAA+ orientation for subunits A and F at the seam (0.4–0.8 Å), indicating minimal conformational variability within the hexamer (Figure 2D). The orientation of the HBD relative to the AAA+ domain appears to be stabilized by intramolecular salt bridges between D252 and two lysine residues, K95 and K99 (Figure 2E).

### Structural basis for ATP binding and hydrolysis

The catalytic site of SF3 ATPases is formed at the interface between two adjacent subunits. One subunit provides the cis-acting Walker A, Walker B, and motif C residues, while the neighboring subunit contributes the trans-acting arginine finger. In our NS3 cryo-EM reconstruction, clear density corresponding to ATPγS is visible in all six nucleotide-binding sites, although it is less well resolved in protomer F, which lies at the seam of the hexamer (Figure 3A–F). At each site, the triphosphate moiety of ATPγS is coordinated by the P-loop, with the β-phosphate forming interactions with the backbone amides of G165, G167, and K168 (Walker A). The lysine side chain is positioned to interact with both the β- and γ-phosphates. In five of the six sites, resolved to ∼2 Å, we observed hexavalent coordination of a Mg²⁺ ion by the β- and γ-phosphates, the side chain of T169, and three water molecules. Solvent molecules in this primary hydration shell form hydrogen bonds with D213 (Walker B) and D231 (part of pore loop 2). A water molecule located between N259 and D213 in these five sites is likely positioned for nucleophilic attack on the γ-phosphate (Figure 3A-D).

**Figure 3:**
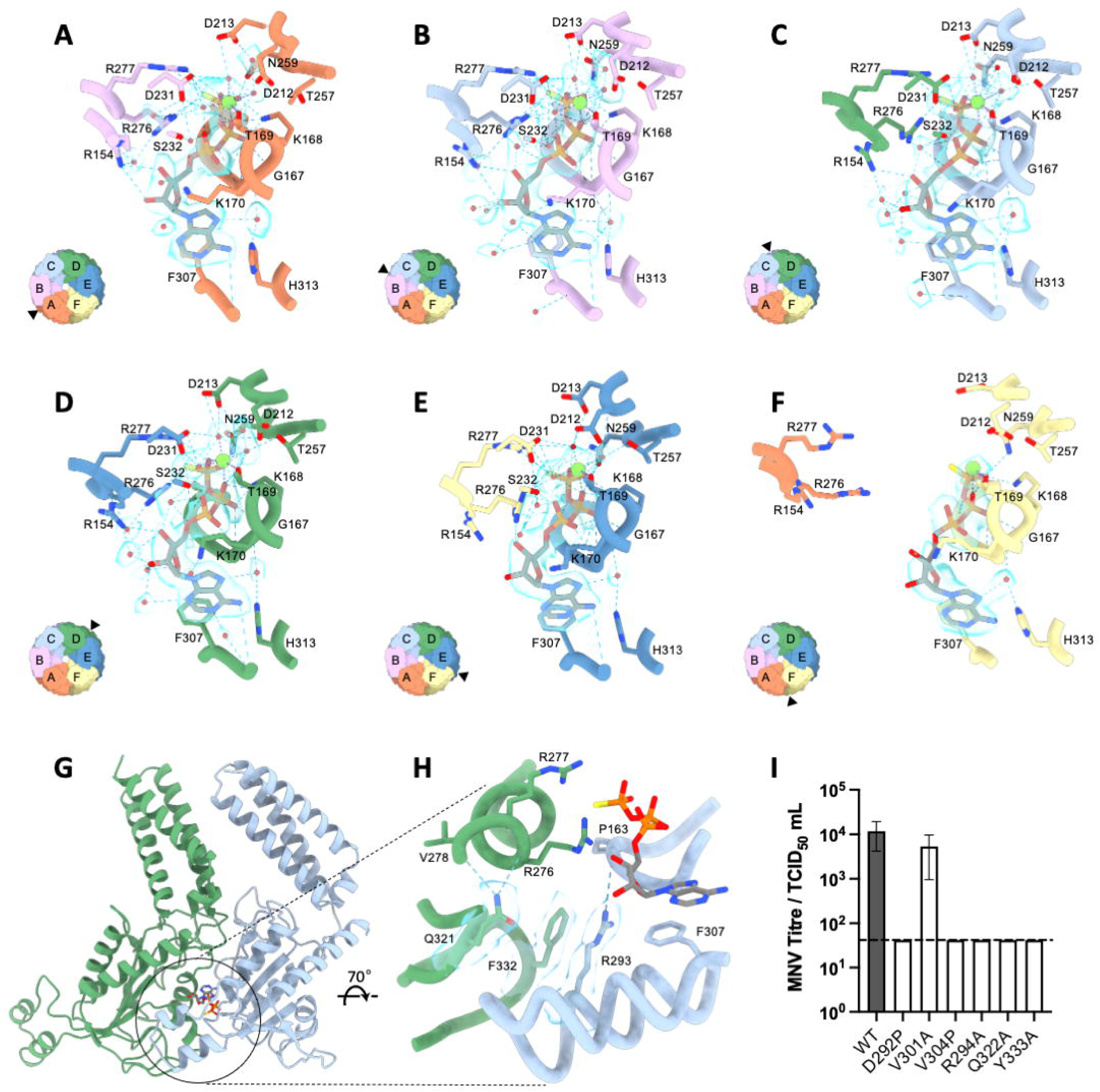
Molecular basis for ATP binding and hydrolysis. (A-F) Zoomed-in view of each of the six ATP binding sites in the NS3 hexamer. The inset panel shows the location relative to the overall structure. (G) Cartoon representation of subunits C and D. (H) Zoomed-in view of the ATP binding site with key residues shown as sticks and their EM density colored blue. (I) WT and mutant MNV were titrated by TCID50 assay on BV-2 cells (n = 3 ± SEM). The dashed line denotes the limit of detection for TCID50 assays.

Subunits A–E show the γ-phosphate interacting with R276 and R277 (arginine finger), as well as with N259 (motif C), supporting their classification as ATP-bound. In contrast, subunit F adopts an apo-like conformation, as the arginine finger is displaced across the seam and does not engage the bound nucleotide (Figure 3F). Additional stabilizing interactions in the ATP-bound subunits include a hydrogen bond between the ribose 3′ hydroxyl of ATPγS and R154. At the base recognition level, the adenine ring is sandwiched between K170 and F307, stabilized through cation–π and π–π stacking interactions. A water molecule is also observed forming hydrogen bonds with the ATPγS imidazole ring, the backbone carbonyl of G167, and the side chain of H313. The nucleotide-binding pocket is further reinforced by a cation–π interaction between R293, located in a helix–turn–helix motif, and F332 in the neighboring protomer. Upstream, Q321 engages in hydrogen bonds with the backbone atoms of V278 and R276, helping to stabilize the helix that presents the arginine finger (Figure 3G–H).

Due to the unavailability of infectious clones for HuNoV, to validate the functional relevance of these residues, we introduced corresponding mutations into the NS3 protein of murine norovirus (MNV) and assessed viral infectivity (Supplementary Figure 9). Mutations that disrupted the cation–π interaction or destabilized the helix– turn–helix motif resulted in complete loss of infectivity, consistent with the structural role these residues play in catalysis (Figure 3I).

### Architecture of the NS3 pore loops

Within our NS3 structure, we identified two pore loops, both of which were resolved at approximately 2–3 Å resolution in our cryo-EM reconstruction. These loops are located beneath a solvent-accessible chamber formed by the HBDs, with an estimated volume of 19,400 Å³ (Figure 4A). Pore loop 2 defines a central channel approximately 15 Å in diameter, sufficient to accommodate single-stranded RNA, into which the hydrophobic side chain of I242 projects. The two pore loops are stabilized by inter-protomer hydrogen bonding and packing of side chains W200, L237 and F249. At the subunit level, pore loops 1 and 2 are closely associated, forming an average interface area of approximately 420 Å² (Figure 4B). With the exception of subunits A and F, located at the seam of the hexamer, pore loops from adjacent subunits form inter-subunit contacts, averaging ∼360 Å^2^ in interface area. These contacts are mediated by electrostatic interactions between D240 and K245, as well as intermolecular hydrogen bonds involving the side chains of R194 and R241 with the backbone carbonyls of K245 and P245, which collectively help stabilize the central pore architecture (Figure 4C).

**Figure 4:**
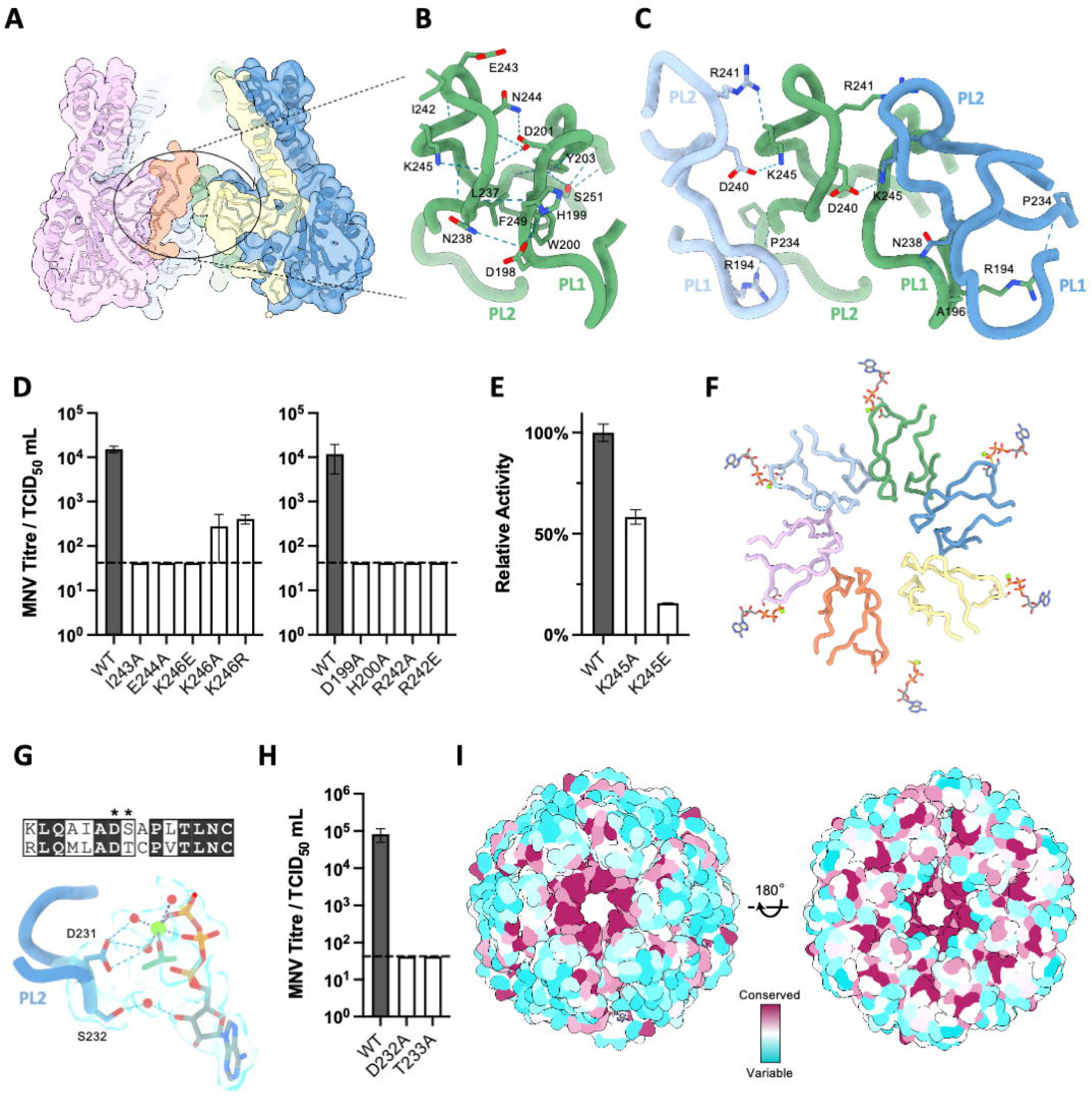
The architecture of the NS3 pore loops. (A) Central slice through the surface representation of the NS3 hexamer. (B) Zoomed-in view of pore loops (PL) 1 and 2 from subunit D, with residues involved in intramolecular interactions shown as sticks. (C) As shown in (B) for subunits C-E, with residues involved in inter-protomer interactions shown as sticks. (D) Infectivity of WT MNV and pore loop mutants (n = 3 ± SEM). (E) ATPase activity of the WT, K245A, and K245E NS3 mutants (n = 2 ± SD). (F) Cartoon representation of the NS3 pore loops, with ATPγS shown as sticks. (G) Zoomed-in view of residues 227-234 of protomer E and the ATPγS molecule located in the DE active site. (H) Infectivity of WT MNV compared to the D232A and T233A NS3 mutants (n = 3 ± SEM). (I) Two orthogonal views of the NS3 hexamer shown as a surface representation and colored according to conservation across 46 NS3 sequences. For panel H, a representative result is shown.

To functionally validate the importance of these structural features, we introduced targeted mutations into the MNV NS3 pore loops. In line with our structural observations, most mutations completely abolished viral infectivity. Only upon mutation of K246 to an alanine or arginine, a small amount of infectious virus was recovered (Figure 4D). To further investigate the role of K245, we introduced K245A and K245E mutations into the recombinant HuNoV hex-NS3 construct and assessed their effects on ATPase activity. Both mutations impaired ATP hydrolysis, with the charge-reversing K245E mutation producing a more pronounced defect (Figure 4E).

In five of the six ATP-binding sites, pore loop 2 residues D231 and S232 form solvent-mediated hydrogen bonds with the bound ATPγS molecule (Figure 4F-G). In contrast, in subunit A, positioned at the top of the spiral near the seam, pore loop 2 is disengaged. This suggests it may be poised to sense the incoming ATP molecule in the adjacent subunit. Together, these observations indicate that coordination between the pore loops and the nucleotide-binding sites plays a central role in orchestrating sequential ATP hydrolysis across the hexamer (Supplementary Figure 10). Consistent with this model, mutations in key pore loop residues resulted in a complete loss of infectivity in the MNV system (Figure 4H). Sequence analysis using ConSurf, based on 46 calicivirus NS3 sequences, revealed that these residues are highly conserved, highlighting their functional importance and potential as targets for pan-genotypic antiviral intervention (Figure 4I).

### Comparison of NS3 to SF3 helicases from DNA viruses

Structural comparison of the NS3 complex with SF3 helicases from single- and double-stranded DNA viruses revealed several notable differences. In NS3, HBDs, which form the walls of the chamber above the AAA+ pore loops, exhibit minimal or no interactions with one another (Figure 5A). In contrast, the corresponding regions in previously characterized structures of the Rep and E1 SF3 helicases form a tightly packed collar that remains planar during the translocation of single-stranded DNA (Figure 5B– C)^19,20^. In these DNA helicases, the collar not only stabilizes oligomeric assembly but also serves as a mechanical anchor point for the conformational changes of the AAA+ domains^19,20^.

**Figure 5:**
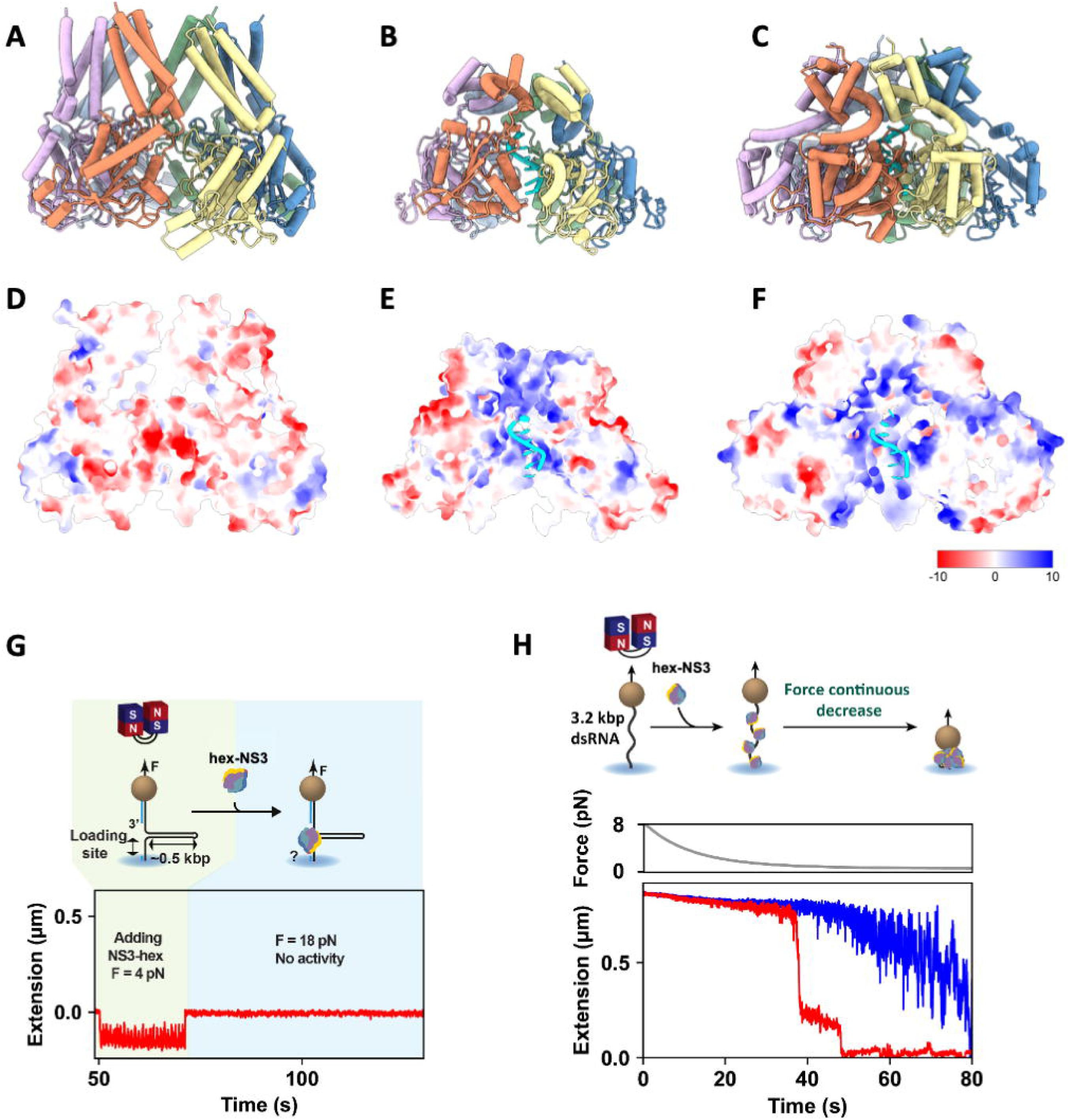
Structural and functional comparison of HuNoV NS3 to known helicases. Cartoon representation of the (A) NS3 hexamer, (B) porcine circovirus 2 rep protein bound to single-stranded DNA (PDB ID: 7LAR)^20^, and (C) papillomavirus E1 hexameric helicase with ssDNA (PDB ID: 2GXA)^19^. (D) Central slice through the NS3 hexamer shown as a surface representation and colored according to electrostatic surface potential. (E) As shown in (D) for porcine circovirus 2. (F) As shown in (D) for papillomavirus E1. (G) Description of the magnetic tweezers assay to probe the helicase activity of hex-NS3. (H) Description of the magnetic tweezers assay to monitor dsRNA compaction by hex-NS3 when decreasing the force continuously. dsRNA tether compaction trace without (blue) and with (red) represents without and with hex-NS3. Panels G and H show representative traces from n = 246 and n = 35 individual bead measurements, respectively. Each assay was performed once, and all measurements were acquired from that single experiment.

By contrast, in NS3, the HBDs and AAA+ domains adopt a shared right-handed helical configuration, suggesting they function synchronously as a unified motor. Another distinguishing feature of NS3 is the pronounced negative electrostatic potential of its central pore (Figure 5D). This contrasts with the positively charged central channels of Rep and E1 (Figure 5E–F)^19,20^, which are electrostatically well-suited to engage the phosphate backbone of DNA. While the negative charge would be incompatible with stable nucleic acid binding in classical helicase function, it is reminiscent of the electrostatic environment observed in other viral nucleic acid translocation systems, where transient interactions may facilitate efficient translocation without stalling ^31–33^.

To investigate the RNA unwinding activity of our hex-NS3 construct, we employed a previously established magnetic tweezers assay. In this setup, magnetic beads were tethered to the surface of a flow chamber using an RNA hairpin construct (Figure 5G and Supplementary Figure 11A). Upon introducing 2.3 µM hex-NS3 and 2 mM ATP into the chamber, we monitored changes in tether extension as a readout for hairpin unwinding. Despite applying a high assisting force, we did not observe any helicase activity from hex-NS3.

We next examined the interaction of hex-NS3 with single-stranded RNA. Upon introducing 1 µM hex-NS3 in the absence of ATP, we observed a delay in RNA hairpin refolding, consistent with engagement of the ssRNA segment (Supplementary Figure 11B). A less pronounced effect was also observed with the MBP-hex negative control, likely due to nonspecific effects (Supplementary Figure 11C).

We then investigated the double-stranded (ds) RNA binding properties of hex-NS3. In this case, magnetic beads were tethered using a ∼3.2 kbp dsRNA construct (Figure 5H and Supplementary Figure 11D). Following the addition of hex-NS3, we gradually reduced the applied force from 8 pN to 0.1 pN. At a critical low force of 1.3 ± 0.2 pN, we observed a sudden compaction of the dsRNA, suggestive of thermally activated long distance interactions. No such compaction was observed when hex-NS3 was replaced with MBP-hex (Figure 5H and Supplementary Figure 11E).

These findings suggest that although our enzymatically active hex-NS3 construct, lacking the authentic NTD, is unable to unwind RNA duplexes, it retains distinct RNA-binding and remodeling properties.

### An integrative model of the full-length NS3 hexamer

As our hex-NS3 construct lacks the NTD, the architecture of the full-length, membrane-associated complex remained unresolved. During the preparation of this manuscript, AlphaFold3 became available^34^, enabling new structure predictions of the full-length hexameric assembly. As with AlphaFold2, the AAA+ ATPase domain and the HBD were predicted with high confidence (Supplementary Figure 12). However, the quaternary structure deviated from the asymmetric configuration observed in our cryo-EM reconstruction (Figure 2A). AlphaFold3 also failed to recapitulate the helical offset and domain staggering within the AAA+ and HBD tiers, instead predicting a more symmetric assembly than seen experimentally. The NTD, by contrast, was initially predicted with low confidence, with most residues assigned pLDDT scores below 50 (Supplementary Figure 12A and C).

Given that the NS3 N-terminus is reported to interact with cellular membranes^15^, we hypothesized that including fatty acid molecules in the input could improve the modeling by mimicking a lipid bilayer environment. Indeed, supplementing the prediction with 50 oleic acid molecules resulted in markedly improved confidence for the NTD, with pLDDT values ranging from 70 to 100 for both HuNoV and MNV (Figure 6A and Supplementary Figure 12B and D).

**Figure 6:**
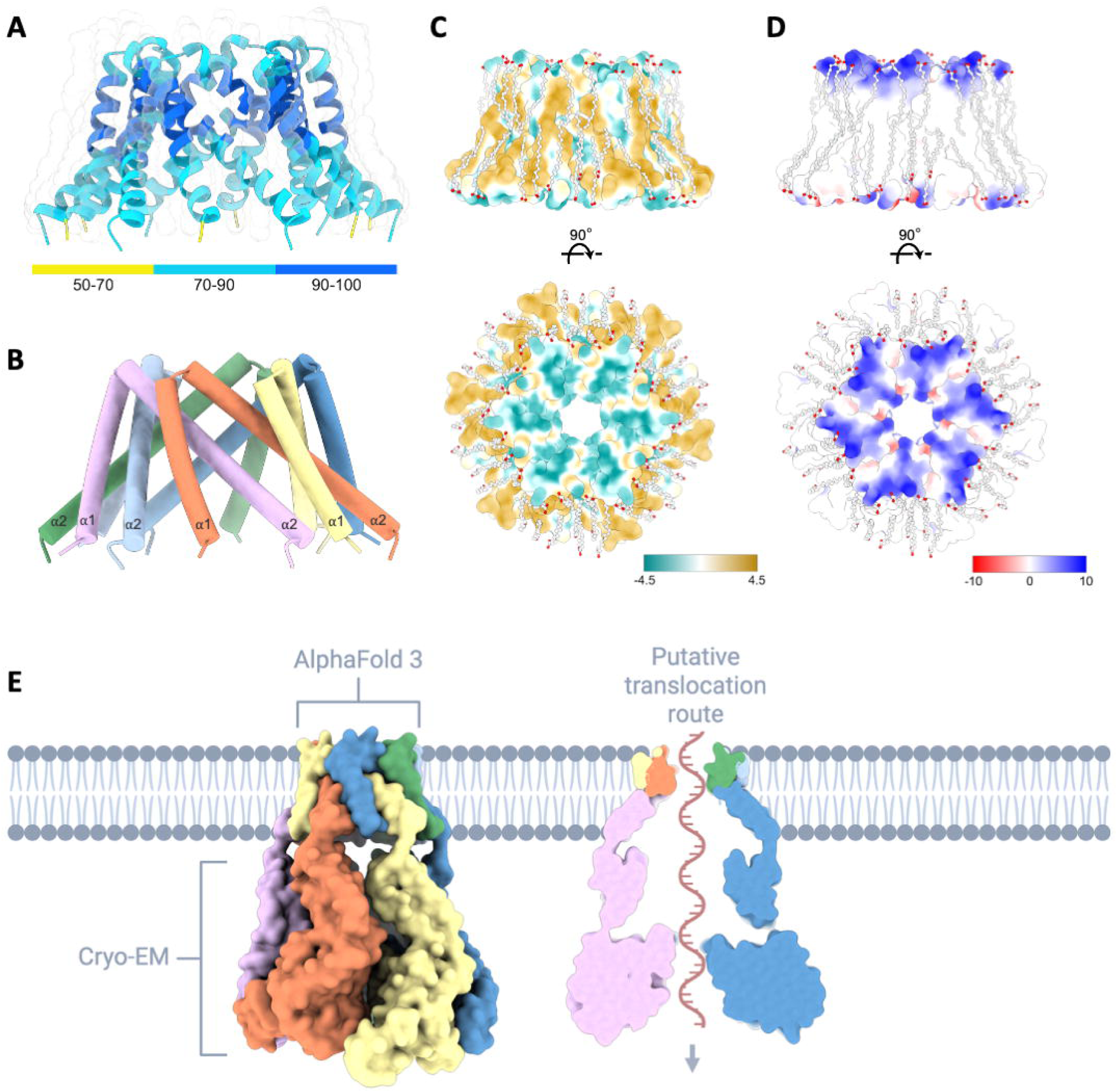
Proposed model for the full-length NS3 complex. (A) Cartoon representation of residues 1–61 from the AlphaFold3 prediction of the full-length HuNoV NS3 hexamer in the presence of 50 oleic acid molecules. The model is colored by confidence score (predicted local distance difference test, pLDDT), with blue indicating high-confidence regions and yellow indicating low-confidence regions. (B) Cylindrical representation of the model shown in (A), with the two transmembrane helices labeled and colored by chain. (C) Surface representation of the NS3 transmembrane pore, colored according to hydrophobicity using the Kyte-Doolittle scale, shown in two orthogonal views. (D) As in (C) but colored by electrostatic surface potential. (E) Surface representation of an integrative model of the full-length NS3 complex embedded in a membrane bilayer (left), and a central cross-section (right) highlighting a putative RNA translocation pathway. Partially created with BioRender.com.

This model suggests that the HuNoV NS3 NTD comprises two transmembrane helices (α1 and α2). Within the hexameric context, these helices intercalate to form a funnel-shaped transmembrane channel (Figure 6B). The height of this region is ∼40 Å, consistent with the thickness of a typical lipid bilayer. The outer surface of the NTD is hydrophobic, while both the cytosolic and luminal surfaces show positive electrostatic potential, features consistent with membrane integration and lipid headgroup interaction. The channel’s interior, with a diameter of ∼15 Å, is positively charged (Figure 6D), potentially favoring interactions with the negatively charged backbone of RNA.

To test the hypothesis that NS3 forms a continuous conduit for RNA, we repeated the AlphaFold3 prediction with the addition of a 50-nucleotide polyadenylate (polyA) RNA strand. Notably, the resulting model positioned the RNA along the central axis of the complex, running uninterrupted through the N-terminal transmembrane channel, the helical bundle, and the AAA+ motor domain (Supplementary Figure 13A). This configuration supports the idea that NS3 can accommodate an extended RNA substrate and suggests a continuous RNA path through the hexameric assembly, consistent with a threading or translocation mechanism.

To extend these observations to related proteins, we performed similar AlphaFold3 predictions with the full-length 2C protein of Coxsackievirus B3. The resulting model closely resembled the NS3 prediction, also featuring a transmembrane channel aligned with a central RNA conduit that traverses the entire complex (Supplementary Figure 13B). These findings are consistent with a conserved architectural, and potentially mechanistic, theme among SF3 ATPases from viruses in the order *Picornavirales*.

To generate a structural model of the full-length, membrane-embedded NS3 hexamer, we integrated our cryo-EM reconstruction of the AAA+ and HBD regions with the AlphaFold3-derived NTD prediction (Figure 6E). This model suggests that NS3 associates with membranes via its N-terminal transmembrane helices and employs a coordinated ATPase cycle to translocate RNA away from the membrane in a hand-over-hand fashion (Supplementary Movie 1).

## DISCUSSION

Despite the clinical and economic impact of human noroviruses, many aspects of their replication cycle remain poorly understood^1,35^. The non-structural protein NS3 has been implicated in several key processes during infection, including a putative role as an RNA chaperone or helicase^12,13^. However, until now, the oligomeric structure of NS3, or of any SF3 ATPase from an RNA virus, had not been resolved.

Replication of norovirus RNA requires the resolution of stable secondary structures and double-stranded RNA intermediates, meaning RNA remodeling is essential for efficient genome synthesis^36^. Although NS3 is widely proposed to support this process, its specific role as an ATP-dependent helicase remains unclear. Both NS3 and its picornaviral homolog 2C exhibit ATPase activity and bind RNA^37–39^, yet reports of their RNA unwinding capabilities have been inconsistent. Some studies have demonstrated ATP-dependent helicase activity, including bidirectional activity for NS3 and 3′ to 5′ activity for 2C^12,40^. Other studies failed to detect unwinding activity, reporting only an RNA-stimulated ATPase function or ATP-independent RNA chaperoning activity^13,37^.

Our soluble NS3 construct lacks the native N-terminal domain, which we replaced with a heterologous hexamerization motif to promote oligomerization. The native NTD is predicted to form a positively charged, membrane-embedded pore, and is likely critical for RNA engagement and directional translocation. Its absence may explain the lack of processive RNA unwinding observed in our assays. These findings may help reconcile prior evidence that full-length NS3 and eukaryotic expression systems are required for helicase activity^12,13^. This dependence may stem from a requirement for host-derived lipids to support correct NTD folding.

Consistent with its proposed role in replication, NS3 has also been shown to colocalize with double-stranded RNA and NS7 at replication organelles^14,41^, further reinforcing its proximity to active sites of genome synthesis. Moreover, NS3 has been shown to stimulate the RNA synthesis activity of the viral polymerase NS7 *in vitro,* suggesting a direct functional interplay between these replication proteins^12^. Our single-molecule magnetic tweezer experiments support a role for the NS3 cytosolic domains in binding both ssRNA and dsRNA, which may reflect either a translocase function or a post-translocation role. Notably, full-length vesicle-associated 2C was recently shown to preferentially retain dsRNA at membranes, suggesting a conserved mechanism for RNA handling at replication organelles^39^.

In addition to its proposed roles in replication, NS3 has been implicated in several other cellular processes, including the induction of apoptosis and the remodeling of host cell membranes^14–16^. Notably, our structural analysis reveals that the HBD of NS3 shares homology with a BAG-family co-chaperone^30^. BAG-domain proteins are known to regulate apoptosis and proteostasis by modulating the activity of HSP70 chaperones^42^. Prior studies have shown that the NS3 HBD is essential for virus-induced apoptosis^16^, raising the possibility that structural mimicry of BAG proteins may contribute to this activity. Supporting a possible role in chaperone modulation, both HSP70 and HSP90 have been implicated as pro-viral in the MNV replication cycle^43^.

With respect to membrane interactions, NS3 has also been shown to associate with various host membranes, including the endoplasmic reticulum, mitochondria and lipid-rich bodies, underscoring its multifunctional role during infection. Overexpression of NS3 induces the formation of cholesterol-rich, vesicle-like structures, suggesting a role in membrane remodeling^14,15,44^. These functions rely on the membrane-binding NTD. Our AlphaFold3-based model of this region, generated in the presence of fatty acids, may explain previous genetic findings. For example, a single amino acid substitution in the interior of the predicted transmembrane pore in MNV was shown to enhance infectivity, underscoring the functional importance of this channel^45^. Another recent study proposed that the NS3 NTD resembles the MLKL pore-forming domain. While our high-confidence model strongly suggests an alternative architecture, this is compatible with the author’s functional observations of lipid binding and membrane permeabilization through the formation of a size-selective pore^44^.

The alignment of the transmembrane domain, central helical bundle, and AAA+ motor domain forms a continuous conduit that may allow RNA to be threaded across replication organelle membranes. The AAA+ pore in our model exhibits a negative electrostatic potential, similar to single- and double-stranded translocation systems observed in other viruses^31–33^, which could facilitate RNA translocation by reducing nonspecific interactions. Our integrative structure of the full-length NS3 complex may have important functional implications. If NS3 indeed facilitates RNA translocation across membranes, it implies that genome synthesis may occur within a protected, membrane-bound compartment from which nascent RNA is exported for either translation or encapsidation. While such a luminal replication site has not yet been definitively identified, tomography studies of MNV and enterovirus-infected cells have identified single- and double-membrane compartments to accumulate in infected cells. Notably, the single-membrane structures observed in these studies could, in principle, support such a topological arrangement and have been proposed as the primary site of replication in the case of poliovirus^46–49^.

Based on the conserved polarity of substrate translocation by SF3 helicases, RNA is likely transported through the channel in a 3′ to 5′ direction. Our NS3 complex diverges from classical SF3 architectures by replacing the static oligomerization domain with a symmetry-matched spiral ring coordinated with the ATPase motor^19,50^ In contrast to the planar HBD tier seen in DNA helicases^2^, in NS3, the equivalent function is likely fulfilled by the transmembrane domain, which may act as a mechanical anchor during ATP-driven RNA translocation. These findings imply that NS3 must be capable of unfolding and translocating the VPg primer covalently attached to the 5′ end of the genome^35^, suggesting that NS3 may function as a dual RNA-protein translocation system.

Our findings not only suggest a new model for norovirus replication but would also reconcile the strategies employed by picornaviruses with those of other positive-strand RNA viruses that remodel host membranes to spatially separate replication from downstream steps^51,52^. It raises questions as to how NS3 governs the polarity, timing, and specificity of RNA export, and how this might be integrated with downstream capsid assembly^53^. Notably, genetic and biochemical studies have previously suggested direct interactions between the picornaviral 2C protein and capsid components^54^, supporting the idea that RNA replication and encapsidation are spatially and functionally coupled. Furthermore, *in situ* cryo-electron tomography of poliovirus-infected cells has revealed viral capsids connected to replication organelles by a proteinaceous complex of approximately 230 kDa^49^. Based on our model, it is reasonable to speculate that this complex may correspond to the 2C hexamer. It would be particularly interesting to perform similar *in situ* structural studies on MNV infected cells, which could help determine whether NS3 adopts an analogous complex on viral replication organelles.

Finally, our structural framework highlights NS3 as a compelling antiviral target. We and others have previously shown that the related 2C protein in enteroviruses can be inhibited by small molecules such as S-fluoxetine, which binds to the predicted pore of the AAA+ domain and allosterically impairs its function^26,55^. Given the structural homology between 2C and NS3, including conservation of the central channel architecture, antiviral compounds could be developed to target norovirus NS3. If the export of newly synthesized genomic RNA is indeed essential for efficient replication, then inhibitors that block RNA translocation through this channel would be expected to disrupt genome synthesis and packaging. This could help explain why such compounds have previously emerged as inhibitors of these processes^55–57^. The high-resolution details of the NS3 ATP-binding pocket and central pore provided here offer a strong foundation for structure-guided antiviral discovery.

In summary, our cryo-EM structure of norovirus NS3, supported by an integrative full-length model, positions this complex as the first example of an SF3 protein functioning as a membrane-spanning RNA translocase, a role distinct from its better characterized DNA virus counterparts^2^. In contrast to the large crown-like replication complexes formed by other positive-strand RNA viruses such as coronaviruses, alphaviruses and nodaviruses^52^, the compact hexameric architecture of NS3 suggests a streamlined mechanism for RNA export across replication organelle membranes. These findings raise the possibility that such minimal, membrane-anchored ATPases may represent an alternative strategy for spatially coupling genome synthesis and packaging, one that could be conserved among the order *Picornavirales*. While further studies will be required to confirm the functional implications of this architecture, our model lays a foundation for future mechanistic investigations and may inform the development of targeted antiviral strategies.

## METHODS

### Cells and reagents

BHK-21 cells (obtained from ATCC) and BV-2 cells (gifted by Ian Goodfellow, University of Cambridge) for MNV experiments were maintained as described previously^58^.

Adenosine 5′-O-(3-Thiotriphosphate) (ATPγS) (Calbiochem [Sigma-Aldrich]) was purchased as a 100 mM solution in water.

### Cloning and construct design

The codon sequence of human norovirus (HuNoV) GI (NC_001959.2) NS3 (amino acids 62-363) optimized for bacterial expression (GenScript) was cloned into the expression vector pET-28b using primer-facilitated ligation. At the N-terminus, a HRV 3C protease-cleavable sequence introduces a Strep-tag II, a His_6_-tag, and MBP-tag to facilitate expression and purification. For the hexameric construct, a 32-residue long codon-optimized cc-hex sequence^29^ followed the 3C protease cleavage sequence at the 5’ end of the NS3 coding sequence (Figure 1A and Supplementary Figure 1B). As a control protein, we used the MBP-hex construct without NS3 sequence. Mutations were introduced via site-directed mutagenesis using Q5® High-Fidelity DNA polymerase (New England Biolabs). All oligonucleotides were purchased from IDT.

The plasmid, pT7-MNV*, containing the infectious clone sequence from MNV-1 strain CW1P3^59^ under control of T7 promoter was used for virus recovery. To exchange NS3 amino acids, standard two-step overlapping PCR mutagenesis was used with pT7-MNV* plasmid as template, exchanging a ApaI to Xhol fragment. The pcDNA3 based expression plasmids used for coupled i*n vitro* transcription and translation assays expressing MNV ORF1 has been previously described^60^.

The sequences of all plasmids were verified by DNA sanger sequencing (Macrogen). The sequences of the primers used in this study are available upon request.

### Protein expression and purification

NS3 proteins were expressed in *E.coli* Rosetta 2 (DE3) (Sigma-Aldrich). Cells were grown in Terrific Broth (TB) medium containing kanamycin (50 µg/mL) and chloramphenicol (34 µg/mL) at 37°C shaking at 180 rpm until an OD_600_ between 0.5-0.8 was reached. After induction of protein expression with 0.5 mM isopropyl β-D-thiogalactopyranoside (IPTG), cells were cultured at 18°C overnight, shaking. Cells were harvested by centrifugation at 5,000g and the cell pellet was stored at -20°C if the purification was resumed later. The pellet was resuspended in lysis buffer (50 mM Tris pH 8.0; 300 mM NaCl; 10 mM imidazole; 1 mM MgCl_2_; 0.1% Triton X-100; 5% glycerol; 0.25 mg/ml lysozyme; 10 µg/ml DNase I; and 1 tablet complete EDTA-free protease inhibitor cocktail per 10 ml buffer) and the samples were sonicated on ice with 8-10 cycles of 30-second sonication and a 15-second pause (Sonics & Materials Inc.). The soluble phase was separated by centrifugation at 30,000g for 45 min at 4°C. The 0.45µm filtered supernatant was applied for immobilized metal affinity chromatography (IMAC) onto a HisTrap HP column (Cytiva) equilibrated with binding buffer (50 mM Tris pH 8.0, 300 mM NaCl, and 25 mM imidazole) using an Äkta go system (Cytiva) with the UNICORN software version 7.6 (Cytiva) at 4°C and isocratically eluted with 500 mM imidazole.

To obtain cleaved constructs, the samples were incubated with HRV type 14 3C protease (Sigma-Aldrich) overnight at 4°C, while dialyzing against above-mentioned binding buffer containing 1 mM dithiothreitol (DTT). A second IMAC step captured the His-tagged 3C protease, the cleaved Strep-His-MBP-tag, and any uncleaved constructs, while the flow-through containing our protein of interest was further concentrated and applied for size exclusion chromatography, using a Superdex 200 Increase 10/300 GL column (Cytiva). The cleaved hexamers were eluted in cryo-EM buffer (25 mM Tris pH 8.0, 300 mM NaCl, and 1 mM MgCl_2_) and the cleaved monomers in storage buffer (10 mM HEPES pH 7.5, 150 mM NaCl, 10 mM MgCl_2_).

The uncleaved constructs were separated by SEC directly after IMAC, using a Superose 6 Increase 10/300 GL column (Cytiva) for hexameric constructs and a Superdex 200 Increase 10/300 GL column (Cytiva) for constructs lacking the cc-hex domain. The samples were eluted with storage buffer.

The purity of all protein samples was confirmed by SDS-PAGE, using denaturing 4x NuPAGE LDS sample buffer (Thermo Fisher Scientific). All concentrated protein samples were aliquoted, flash-frozen in liquid nitrogen, and stored at -80°C until further analysis.

### NTPase assay

The release of inorganic phosphate during NS3-mediated hydrolysis of nucleotide triphosphates (NTPs) was measured using the Malachite Green Phosphate Assay Kit (Sigma-Aldrich) for end-point reaction evaluation, as previously reported with minor changes^26^. In brief, 250 nM enzyme in a final volume of 50 µL reactions in ATPase reaction buffer (10 mM HEPES pH 7.5, 25 mM NaCl, 10 mM MgCl_2_, and 1 mM DTT) was first incubated with the depicted ATPγS concentration for 30 min at room temperature (where applicable) before 1 mM NTP (Thermo Fisher Scientific for all except TTP from Sigma-Aldrich) was added to start the reaction. After 60 min incubated at 37°C, the reactions were diluted 20-fold with Milli-Q water (950 µL) to avoid precipitation downstream during the malachite green reaction. 80 µL of diluted samples were added to 20 µL of AB reagent in clear flat-bottom 96-well plates (Corning 3599). After color development of 30 min at room temperature, the absorbances were read at 630 nm on a microplate reader (FLUOstar Omega). A standard curve allowed quantification of the samples.

### Mass photometry

Mass photometry measurements were carried out on a OneMP Mass Photometer (Refeyn Ltd). Glass microscope coverslips (24 mm x 50 mm, Paul Marienfeld GmbH) were sonicated for 10 min in isopropanol (HPLC grade), then Milli-Q water, repeating 3 times, before they were dried using a clean N_2_ stream and a clean CultureWell gasket (Grace Biolabs) was applied. Mass calibration was done using an in-house mix of IgG halfbodies (73 kDa), IgG1 (149 kDa), apoferritin (513 kDa), and IgM (967 kDa). Protein dilutions were carried out immediately before measurements. For the experiments, 12 μL sample buffer was loaded into the well to find focus and 3 μL of purified protein sample was added for measurements. The final concentrations of the samples ranged approximately from 10 to 50 nM. Movies were recorded for 60 sec using the AcquireMP software (Refeyn Ltd) and consequently analyzed using DiscoverMP (Refeyn Ltd, version v2023).

### Native mass-spectrometry

Native MS experiments were carried out on a Thermo Scientific Exactive Plus EMR mass spectrometer with a static spray set-up, using in-house produced gold-coated electrospray borosilicate capillaries. Purified cleaved NS3 and hex-NS3 constructs were buffer-exchanged six times into 150 mM ammonium acetate solution (pH 7.0) using Amicon Ultra 0.5 mL 10 kDa centrifugal filters (Merck Millipore). The concentration was calculated using the A280 extinction measured on a Nanodrop One system (Thermo Scientific) and the theoretical extinction coefficient based on the sequence of the construct in the unreduced form^61^. These were 17.335 M^-1^cm^-1^ for cleaved NS3 and 24.325 M^-1^cm^-1^ for cleaved hex-NS3. Samples were diluted for native MS using the same ammonium acetate solution. Cleaved NS3 was diluted to 10 μM and cleaved hex-NS3 was diluted to 25 μM. Spectra were analyzed and deconvoluted using UniDec version 6.0.4.

### Negative-stain electron microscopy

Cleaved hex-NS3 was diluted to 0.03 mg/ml in ATPase reaction buffer containing 1 mM ATPγS, on ice. Subsequently, 3 μl of the sample was applied to carbon-coated copper grids that had been glow discharged for 30 sec using a Cressington 208 instrument. The sample was allowed to absorb for 30 sec prior to two washes with Milli-Q water and staining with 2% uranyl acetate solution. Grids were allowed to dry in air for ∼5 min and then imaged on a Thermo Scientific Talos L120C transmission electron microscope (TEM) operated at 120 keV, equipped with a CCD camera.

### Sample preparation for cryo-EM

Cleaved hex-NS3 was diluted to 0.35 mg/ml in ATPase reaction buffer containing 1 mM ATPγS, on ice. Approximately 3 µl of the complex was applied to QUANTIFOIL R1.2/1.3 grids overlaid with 2 nm layer of carbon that had been glow-discharged for 30 sec on a GloQube (Quorum) at 20 mW power. The sample was applied at 4 °C and 95% relative humidity inside a Vitrobot Mark IV (Thermo Fisher Scientific). The grids were then blotted for 7 sec with +2 blot force and plunge-frozen in liquid ethane.

### Cryo-EM data acquisition

A single grid was imaged on a Thermo Scientific Glacios Cryo-TEM equipped with a Falcon 4i direct electron detector and Selectris energy filter using EPU 2 acquisition software. A total of 1010 movies were collected at a nominal magnification of 130,000x in EER format at 0.9 Å per pixel with total dose of 50.5 electrons per Å^2^. Defocus targets cycled from −0.75 to −1.5 μm. A summary of all data collection parameters is shown in Supplementary Table 1.

### Single-particle image processing

Patch motion correction and patch CTF estimation were carried out in cryoSPARC^62^. Templates for particle picking were generated from a 20 Å resolution map generated from the NS3 AlphaFold2 coordinates using the UCSF Chimera molmap command^63,64^. A total of 1,229,229 picked particles were extracted in a 76-pixel box. During extraction, particles were Fourier binned by a non-integer value, resulting in a final pixel size of 3.32 Å. A single round of two-dimensional (2D) classification was carried out, after which a subset of 2D classes containing 235,173 particles, corresponding to top and side views of the NS3 complex, were selected. Ab initio reconstruction with four classes generated one well-defined reconstruction of the NS3 hexamer and three junk classes. The resulting ab initio maps were then used as initial models for iterative rounds of heterogeneous refinement, where the particles belonging to the NS3 class were used as the input for subsequent rounds. To avoid missing NS3 particles that may have been removed during initial stringent selection of 2D classes, heterogeneous refinement was carried out on the initial particle stack of 1,229,229 particles. Five rounds of heterogeneous refinement yielded a well-defined reconstruction of the NS3 complex containing 123,643 particles. These were then re-extracted unbinned in a 300-pixel box, resulting in a final pixel size of 0.9 Å. Subsequently, non-uniform refinement was carried out on the extracted particles with C1 symmetry imposed, yielding a reconstruction with a global resolution of 2.9 Å. The gold standard Fourier shell correlation (FSC) criterion (FSC = 0.143) was used for calculating all resolution estimates, and 3D-FSC plots were generated in the 3DFSC Processing Server^65^. To facilitate model building, the globally refined map was filtered by local resolution in cryoSPARC. An overview of the data processing pipeline is shown in Supplementary Figure 5.

### Model building and refinement

An AlphaFold2 model was used as a starting point for modeling the NS3 hexamer^64^. First, the predicted hexameric model (residues 62-363) was fitted into the EM density map using the UCSF Chimera Fit in Map tool^63^. Next, molecular dynamics flexible fitting in Namdinator was employed to account for differences in the quaternary structure of the predicted model and the experimental map^66^. Water molecules, magnesium ions, and ATPγS molecules were added in Coot^67^. Subsequently, iterative rounds of modelling in ISOLDE and refinement in Phenix were carried out to improve non-ideal rotamers, bond angles, and Ramachandran outliers^68,69^. During real-space refinement, secondary restraints were imposed. Phenix ReadySet was used to generate restraints for coordinated Mg^2+^ ions. Model validation was carried out using MolProbity^69,70^. All data collection, image processing, and refinement information can be found in Supplementary Table 1.

### Analysis and visualization

Protein-protein and protein-ligand interactions were analyzed using PDBePISA and LigPlot+^71,72^. The volume of the NS3 HBD chamber was calculated using CASTpFold^73^. Structural homology analysis was performed in Foldseek and DALI^74,75^, and structural phylogenetic analysis was carried out using FoldTree^76^. Figures were generated using UCSF ChimeraX^77^. Structural biology applications used in this project were compiled and configured by SBGrid^78^.

### MNV *in vitro* transcription and virus recovery

MNV plasmids were linearized with *NotI* and phenol/chloroform extracted before being used for *in vitro* transcription using the HiScribe™ T7 ARCA mRNA Kit (New England Biolabs), following the manufacturer’s instructions, as previously described^58^. RNA was purified and concentrated using the RNA Clean and Concentrator Kit (Zymo). RNA/DNA transfection was carried out as previously described^58^.

### TCID_50_ assay

Viral infectivity was determined using a TCID_50_ assay as described previously^79^. Briefly, BV-2 cells were seeded into 96-well plates at 2×10^4^ cells/well and left overnight before infection. TCID_50_ values were calculated according to Spearman and Kärber.

### Statistical analysis

Statistical analysis and data visualization were performed using GraphPad Prism version 10. Biochemical experiments were independently repeated on at least two separate days, with each experiment conducted in technical duplicates or greater. Virus infection experiments were performed in at least three independent biological replicates, each including technical duplicates. No statistical methods were used to predetermine sample size, and no data were excluded from the analysis.

### RNA Hairpin Fabrication for Helicase Activity

The helicase hairpin was constructed following a protocol similar to that used for the RNA hairpin described previously^80^. The key difference lies in the orientation of the docking site: for helicase activity, the docking site was positioned on the 5′-end handle, opposite to the 3′-end docking used for polymerase activity, reflecting the opposite translocation polarities of the helicase and polymerase. The helicase docking site was ∼20 nucleotides long, compared to 25 nucleotides for the polymerase. The loop oligonucleotide in the helicase hairpin was 4 nucleotides long, versus 20 nucleotides in the polymerase construct.

### Double-Stranded RNA Synthesis

The synthesis of ∼3.2 kbp dsRNA followed the protocol described previously^81^. Briefly, DNA templates were amplified from a pBAD plasmid using Phusion polymerase and primers containing a T7 promoter. In vitro transcription was performed using the HiScribe Kit (New England Biolabs) to generate ssRNA. The 5′ ends of the ssRNA fragments were monophosphorylated using 5′-polyphosphatase (Biosearch Technologies). Complementary ssRNA strands were annealed to form dsRNA and ligated to RNA handles (452 bp and 513 bp) containing digoxigenin-UTP and biotin-UTP (Jena Biosciences) using T4 RNA ligase 2 (New England Biolabs).

### Flow Cell Assembly and Surface Functionalization

Flow cells were assembled as described previously^82^. A double layer of Parafilm was sandwiched between two #1 glass coverslips. The top coverslip had two holes for fluid access, while the bottom was coated with 0.1% (w/v) nitrocellulose in amyl acetate. The assembled cell was mounted in a custom holder and flushed with ∼1 mL of phosphate-buffered saline (PBS). Polystyrene beads (3 µm diameter; LB30, Sigma-Aldrich), diluted 1:1000 in PBS, were incubated in the chamber for 3 minutes to allow non-specific adsorption to the surface. After rinsing with PBS, 50 µL of anti-digoxigenin (50 mg/mL in PBS) was introduced and incubated for 30 minutes. Excess antibody was removed with 1 mL of high-salt TE buffer (10 mM Tris, 1 mM EDTA, pH 8.0, 750 mM NaCl, 2 mM sodium azide), followed by 0.5 mL of standard TE buffer (150 mM NaCl). Finally, the surface was passivated with 10 mg/mL bovine serum albumin (BSA) in PBS with 50% glycerol for 30 minutes and rinsed with TE buffer.

### Single-Molecule Helicase Unwinding Experiments

Streptavidin-coated Dynabeads M-270 (20 µL; Thermo Fisher Scientific) were mixed with ∼0.1 ng of RNA hairpin (in 40 µL total volume) and incubated for 5 minutes. Beads were rinsed with ∼2 mL of TE buffer to remove excess RNA and unbound beads. Functional hairpins were identified based on the characteristic extension jump (∼0.6 µm at 30 pN) observed during force ramping, indicative of hairpin opening.

The flow cell was flushed with 0.5 mL reaction buffer (20 mM HEPES, pH 7.0, 25 mM NaCl, 0.5 mM MgCl₂, 1 mM TCEP), followed by 100 µL of a reaction mix containing 2.3 µM cleaved hex-NS3 and 2 mM ATP. Hairpins were held closed at a constant force of 18 pN. Prior to measurements, hairpins were opened and closed twice to ensure tether integrity. The magnet was then fixed at a height corresponding to 18 pN for 20 minutes while helicase activity was monitored. Data were acquired at 58 Hz, with temperature maintained at 25 °C. Mechanical drift correction was performed by subtracting the position of a reference bead from the magnetic bead position and the focus of the objective was fixed in respect of the flow cell surface^83^.

### Single-Molecule dsRNA Compaction Experiments

Dynabeads MyOne streptavidin T1 (10 µL; Thermo Fisher Scientific) were washed in 40 µL of 1× TE buffer, mixed with ∼0.1 ng of 3,149 bp dsRNA, and processed as in the unwinding experiments to remove excess RNA and beads. Tether selection criteria were as described previously^84^.

The flow cell was flushed with 0.5 mL reaction buffer (same composition as above). At 4 pN force, reaction buffer containing either no protein, 4.6 µM hex-NS3, or 5 µM MBP-hex was introduced and incubated for 5 minutes. Force-extension measurements were then performed by gradually raising the magnets at 0.1 mm/s, reducing force from 8 pN to 0.1 pN, as previously described^85^. Experiments were conducted at 25 °C, with data acquired at 58 Hz.

For stepwise force-extension experiments, 20 µL of M-270 beads were mixed with ∼0.1 ng of dsRNA in a total volume of 40 µL. The protocol up to tether selection was identical. After flushing with 0.5 mL of reaction buffer, 50 µL of 4.6 µM cleaved hex-NS3 was introduced at 10 pN and incubated for 2 minutes. The magnet was then moved stepwise to forces of 5, 3, 1, 0.3, and 0.1 pN, with a 4-minute dwell at each step. Traces were recorded at 58 Hz.

### Compaction Force Analysis

To calculate the average force at which dsRNA compaction occurred in the presence of hex-NS3, the magnet position corresponding to the compaction event was recorded and the applied force was back-calculated^85^. The error represents the standard error of the mean.

## DATA AVAILABILITY

Atomic coordinates have been deposited in the Protein Data Bank under accession code 9R34. The corresponding EM density maps, including the final unsharpened, sharpened, local resolution–filtered maps, half maps, and mask, have been deposited in the Electron Microscopy Data Bank under accession code EMD-53547. Unaligned gain-normalized movies, motion-corrected micrographs, and the final refined particle stacks have been deposited in the Electron Microscopy Public Image Archive under accession code EMPIAR-12758. The integrative model of the full-length NS3 hexamer is available at https://doi.org/10.5281/zenodo.15556769. Requests for reagents should be directed to the corresponding author, D.L.H.

## ACKNOWLEDGEMENTS

D.L.H. is supported by a Dutch Research Council (NWO) Veni grant (VI.Veni.212.102) and by the European Union (ERC, PicAAA, 101164550). J.S. is supported by the Dutch Research Council NWO Gravitation 2013 BOO, Institute for Chemical Immunology (ICI; 024.002.009), and the European Union (ERC, FLAVIR, 101077640). F.J.M.vK. is supported by the European Union (ERC, VIRLUMINOUS, 101053576). Views and opinions expressed are however those of the authors only and do not necessarily reflect those of the European Union or the European Research Council. Neither the European Union nor the granting authority can be held responsible for them. This work was supported by funding to M.R.H. from the Medical Research Council (MR/S007229/1). D.D. is supported by the BaSyC – “Building a Synthetic Cell” Gravitation grant (024.003.019) from the Netherlands Ministry of Education, Culture and Science and the NWO. D.L.H. also acknowledges funding from the Beijerinck Premium of the M.W. Beijerinck Virology Fund, Royal Netherlands Academy of Arts and Sciences (KNAW). We thank members of the Utrecht Virology Lab for valuable feedback during the preparation of this manuscript, and the staff of the Utrecht University Electron Microscopy Centre for their technical assistance.

## AUTHOR CONTRIBUTIONS

D.L.H. conceptualized the project. M.H. and D.L.H. designed the biochemical and cryo-EM experiments, including the design of protein constructs. C.K. and J.S. designed the mass spectrometry experiments. A.D. and D.D. designed the magnetic tweezers experiments. J.M. and M.R.H. designed the virology experiments. M.H., H.H., C.N., and T.D. cloned the protein constructs and performed protein expression and purification. M.H., H.H., C.N., and T.D. conducted NTPase assays. M.H. and D.L.H. performed AlphaFold predictions. M.H. carried out negative-stain EM experiments. I.D. prepared cryo-EM samples and collected data. M.H. and D.L.H. processed cryo-EM data and built and refined the atomic models. C.K. performed mass photometry and mass spectrometry experiments. C.K. and J.S. analyzed the mass spectrometry data. J.M. and M.R.H. generated MNV infectious clones and conducted virology experiments. A.D. performed the magnetic tweezers experiments. A.D. and D.D. analyzed and interpreted the magnetic tweezers data. M.H. and D.L.H. analyzed the structural data. M.H., J.M., C.K., A.D., D.D., J.S., M.R.H., and D.L.H. curated data. M.H., C.K., A.D., J.S., and D.L.H. visualized the data. F.J.M.vK., D.D., J.S., M.R.H., and D.L.H. secured funding. I.D., F.J.M.vK., D.D., J.S., M.R.H., and D.L.H. provided resources including equipment and reagents. F.J.M.vK., D.D., J.S., M.R.H., and D.L.H. supervised the project. D.L.H. coordinated the overall project administration. D.D., J.S., M.R.H., and D.L.H. coordinated and supervised experimental work. M.H. and D.L.H. wrote the initial draft of the manuscript. All authors contributed to the review and editing of subsequent versions.

