## Supplementary information for "Integrative Structure of Norovirus NS3 Suggests a Role in RNA Transport"

**This PDF file includes:**

Figures S1 to S13

Table S1

Caption for Video S1

**Other Supplementary Materials for this manuscript include the following:**

Video S1

**A**

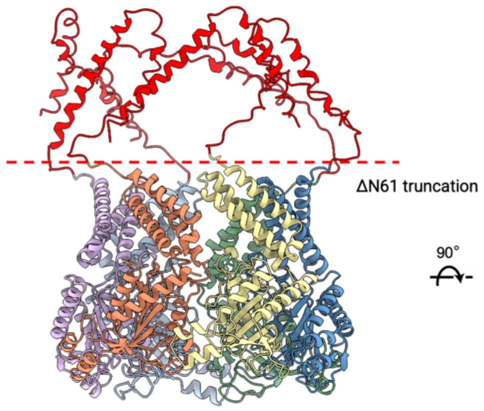

**B**

Strep-tag II, 6xHis

$H_3N^+$-MGSSWSHPQFEKSAHHHHHHMKIEEGKLVIWINGDKGYNGLAEVGKKFEKDTGIKVTVEH

Matlose-binding protein

PDKLEEKFPQVAATGDGPDIIFWAHDRFGGYAQSGLLAEITPDKAFQDKLYPFTWDAVRY

NGKLIAYPIAVEALSLIYNKDLLPNPPKTWEEIPALDKELKAKGKSALMFNLQEPYFTWP

LIAADGGYAFKYENGKYDIKDVGVDNAGAKAGLTFLVDLIKNKHMNADTDYSIAEAAFNK

GETAMTINGPWAWSNIDTSKVNYGVTVLPTFKGQPSKPFVGVLSAGINAASPNKELAKEF

LENYLLTDEGLEAVNKDKPLGAVALKSYEEELAKDPRIAATMENAQKGEIMPNIPQMSAF

Linker, HRV-3C, Linker

WYAVRTAVINAASGRQTVDEALKDAQTGGGGSLEVLFQGPGGGGSGGGGSGELKAIAQEL

cc-hex, Shi linker

KAIAKELKAIAWELKAIAQGAGGSGSYFQSNAGGDQTEKTLKDIESAVIDMEVLSSTSVT

huNoV GI NS3 (62-363)

QLVRDKQSARAYMAILDNEEEKARKLSVRNADPHVVSSTNALISRISMARAALAKAQAEM

TSRMRPVVIMMCGPPGIGKTKAAEHLAKRLANEIRPGGKVGLVPREAVDHWDGYHGEEVM

LWDDYGMTKIQEDCNKLQAIADSAPLTLNCDRIENKGMQFVSDAIVITTNAPGPAPVDFV

NLGPVCRRVDFLVYCTAPEVEHTRKVSPGDTTALKDCFKPDFSHLKMELAPQGGFDNQGN

TPFGKGVMKPTTINRLLIQAVALTMERQDEFQLQ-$COO^-$

**Figure S1: Construct design for hex-NS3. (A)** Two orthogonal views of the AlphaFold2[1] prediction of the full-length NS3 hexamer with the individual protomers shown in different colors. The ΔN61 truncated N-terminus is depicted in red. **(B)** Amino acid sequence of the MBP-hex-NS3 construct, including a heterologous hexamerization domain and linker described previously[2–4]. A human Rhinovirus (HRV) type-14 3C cleavage site separates the StrepII-, His-, and MBP-tag N-terminally from the cc-hex and ΔN61 truncated NS3 of HuNoV GI (NCBI Reference Sequence: NP_056820.1)

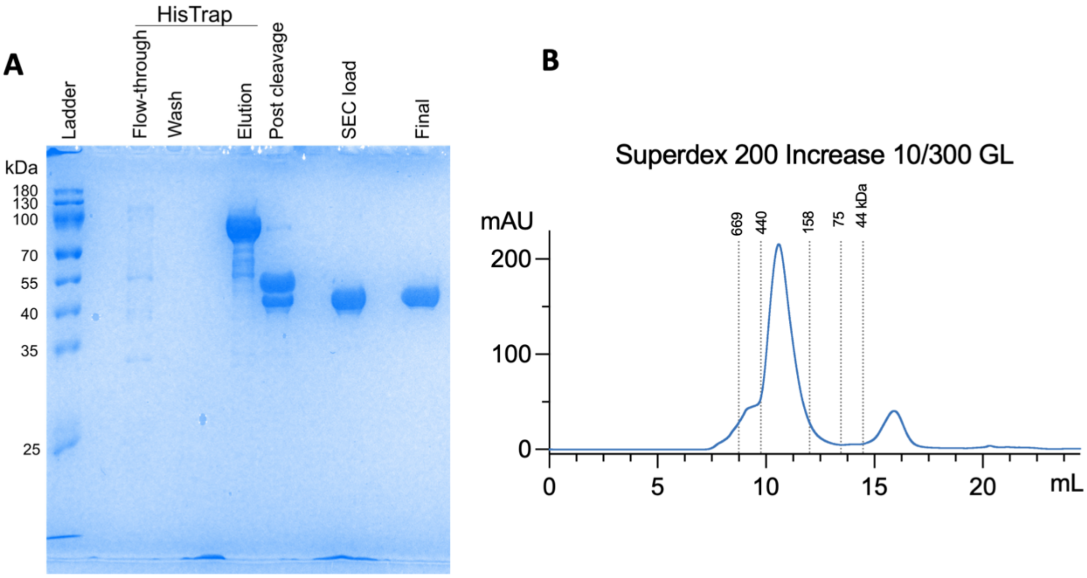

**Figure S2: Purification overview for hex-NS3. (A)** Samples from the individual purification steps are visualized on an SDS-PAGE gel. **(B)** Size-exclusion profile of the cleaved hex-NS3 on a Superdex 200 Increase 10/300 GL column (Cytiva). Calibration markers are indicated with dotted lines and corresponding masses. mAU, milliAbsorbance Unit at 280nm.

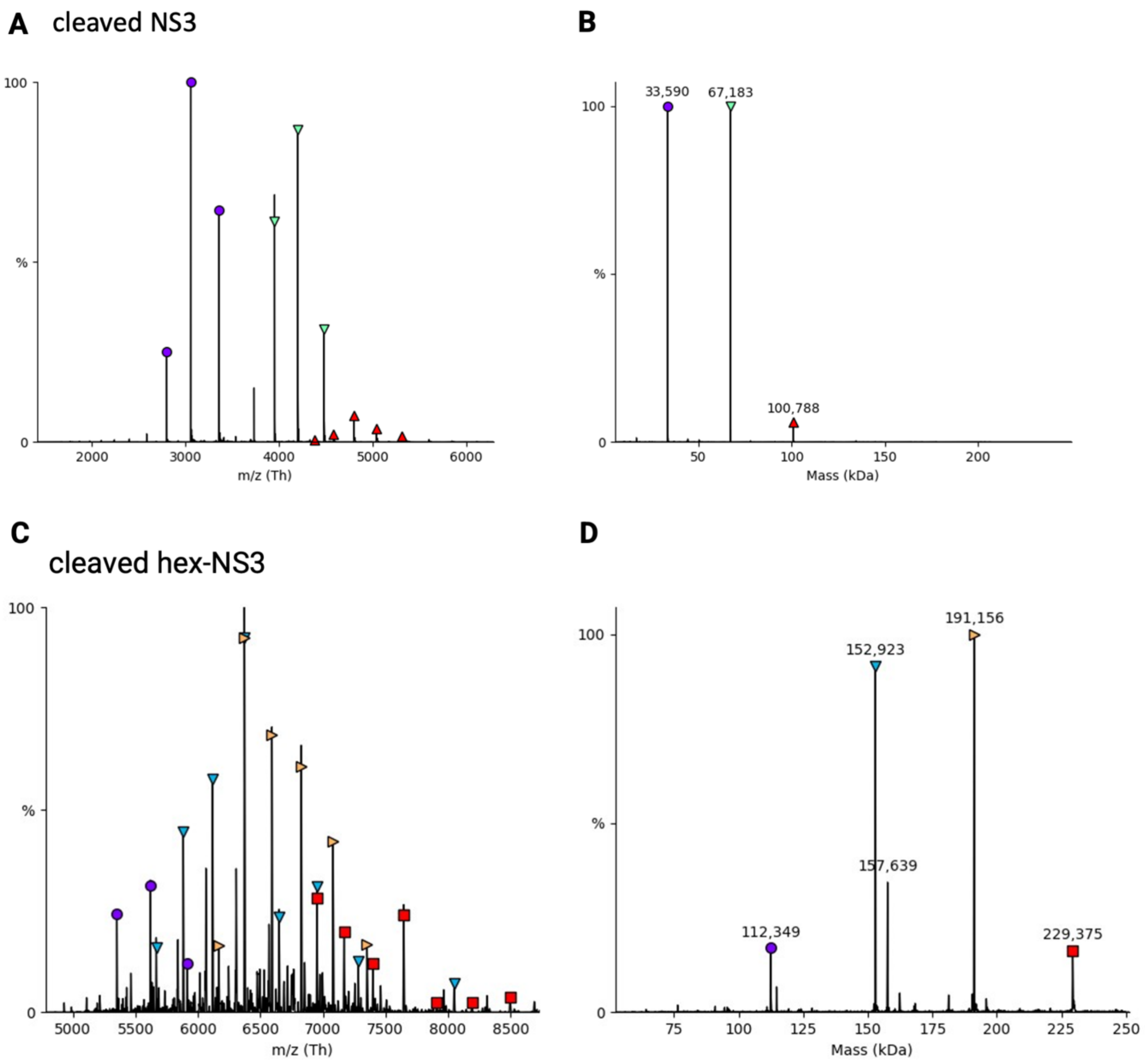

**Figure S3: Native mass-spectrometry analysis of the cleaved NS3 constructs.** **(A)** Native MS spectrum of cleaved NS3 at 10 µM concentration and **(B)** the deconvoluted spectrum thereof. **(C)** Native MS spectrum of cleaved hex-NS3 at 25 µM (based on the protomeric mass), and **(D)** the deconvoluted spectrum thereof.

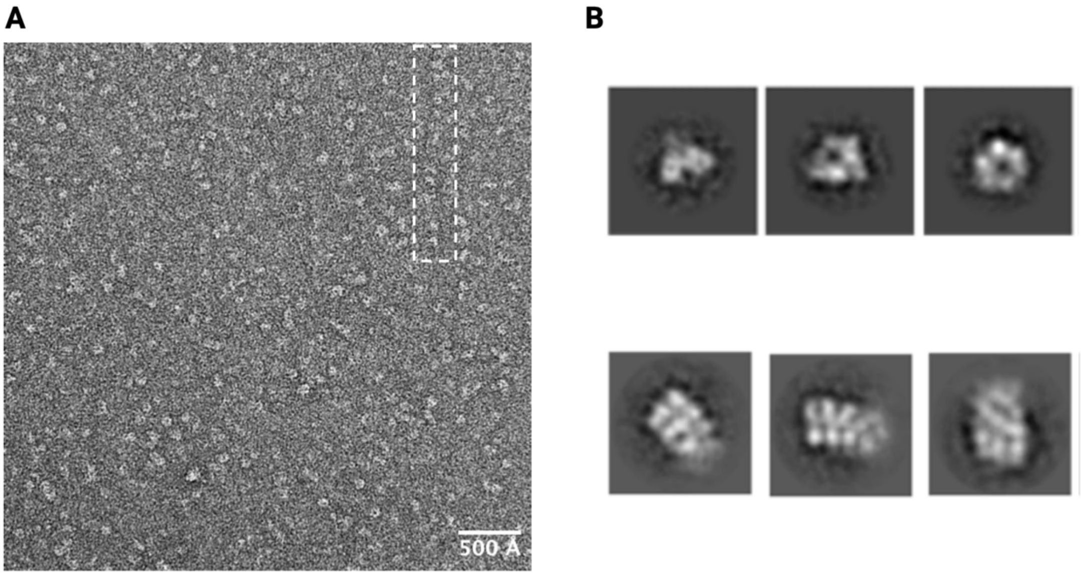

**Figure S4: Negative-stain electron microscopy analysis of hex-NS3. (A)** A representative micrograph of negative-stained cleaved hex-NS3 incubated with ATPγS. The dashed box indicates an example of observed filamentous structures. **(B)** Representative 2D class averages of NS3 hexamers and putative NS3 filaments.

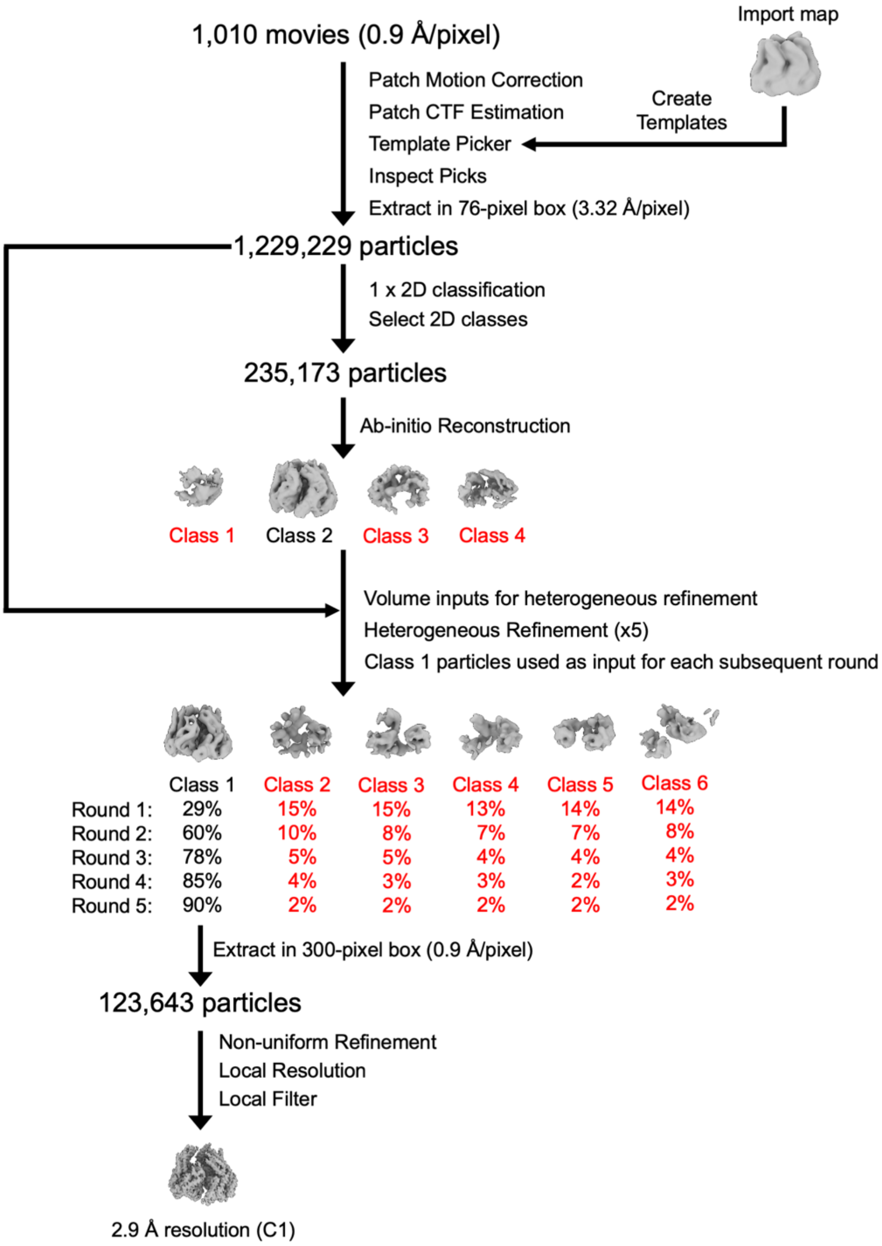

**Figure S5: Single-particle cryo-EM data processing pipeline for hex-NS3 in complex with ATPγS.** Schematic representation of the single-particle cryo-EM data processing pipeline for the hex-NS3 complex. The final reconstruction yielded a 2.9 Å resolution map. Additional details are described in the single-particle image processing section of the materials and methods. CTF: contrast transfer function.

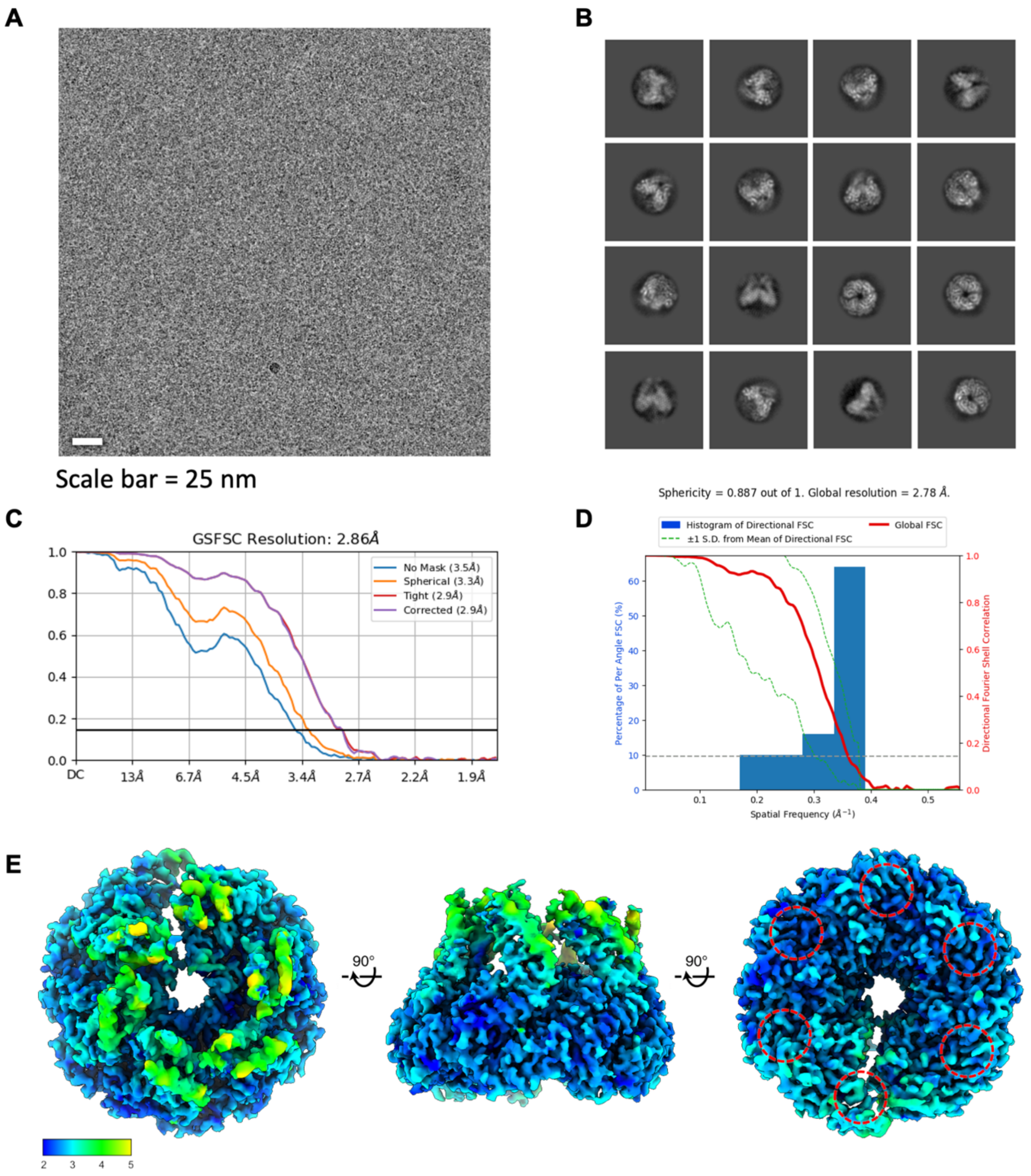

**Figure S6: Single-particle cryo-EM data processing for the hex-NS3 in complex with ATPγS. (A)** Representative motion-corrected electron micrograph of hex-NS3 in complex with ATPγS embedded in vitreous ice. Scale bar = 25 nm. **(B)** Representative reference-free 2D class averages. **(C)** Gold-standard Fourier shell correlation (FSC) curve generated from the independent half maps contributing to the 2.9 Å global resolution density map. **(D)** 3DFSC plot for the hex-NS3 complex[5]. **(E)** Three orthogonal views of the local resolution filtered EM density map for the refined hex-NS3 complex colored according to local resolution which was calculated in CryoSPARC[6]. The locations of the ATPγS binding sites are circled.

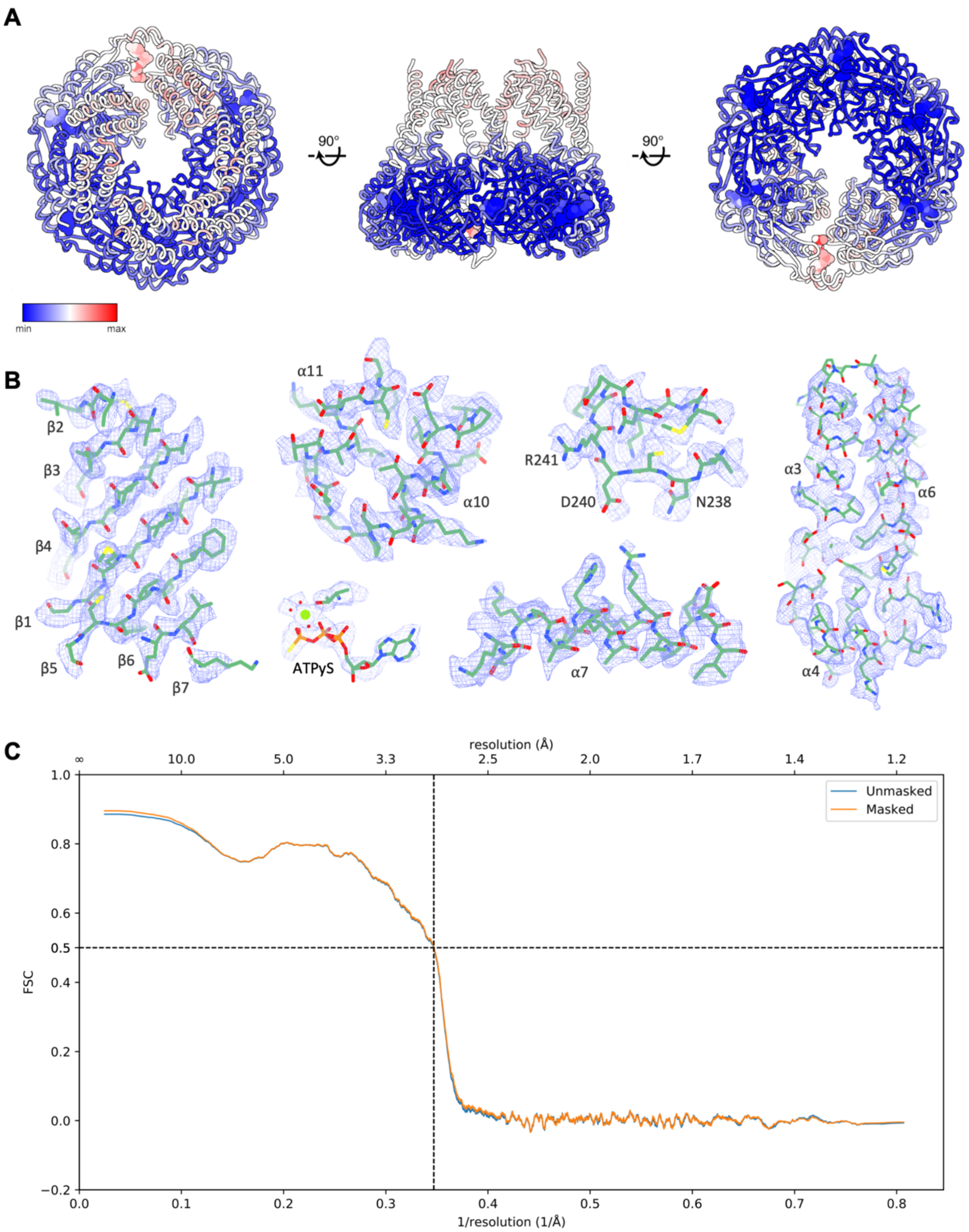

**Figure S7: Atomic modelling of hex-NS3 in complex with ATPγS. (A)** Atomic model of NS3 complex shown as three orthogonal views, with residues colored according to calculated B-factor. **(B)** Representative EM density and fitted atomic model for chain C of the hex-NS3 complex. **(C)** Map versus model FSC curves for the final refined model, generated in Phenix[7].

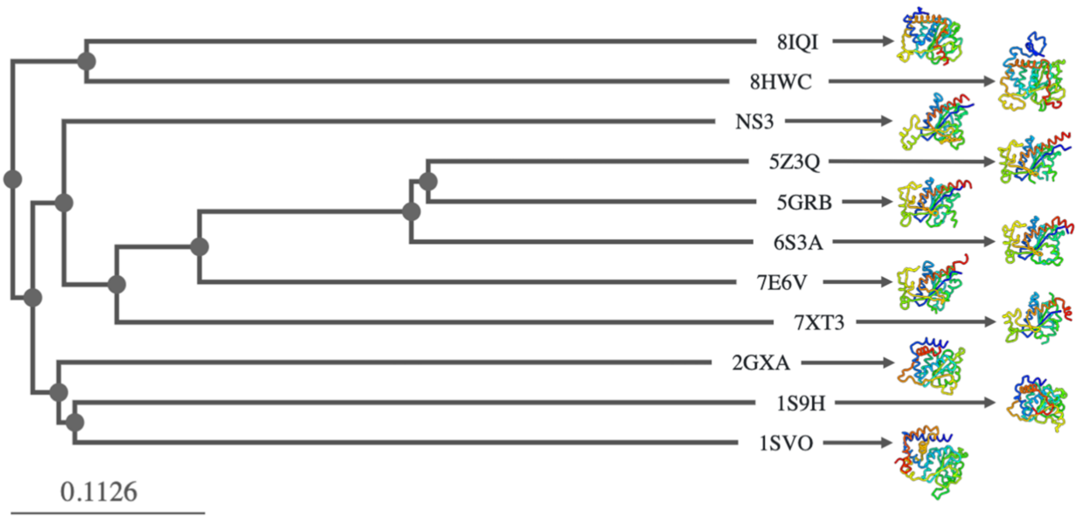

**Figure S8: Structure-based phylogenetic classification of the NS3 AAA+ domain.** Structure-based phylogenetic classification of experimentally determined AAA+ ATPase domains from DNA and RNA viruses calculated with FoldTree[8].

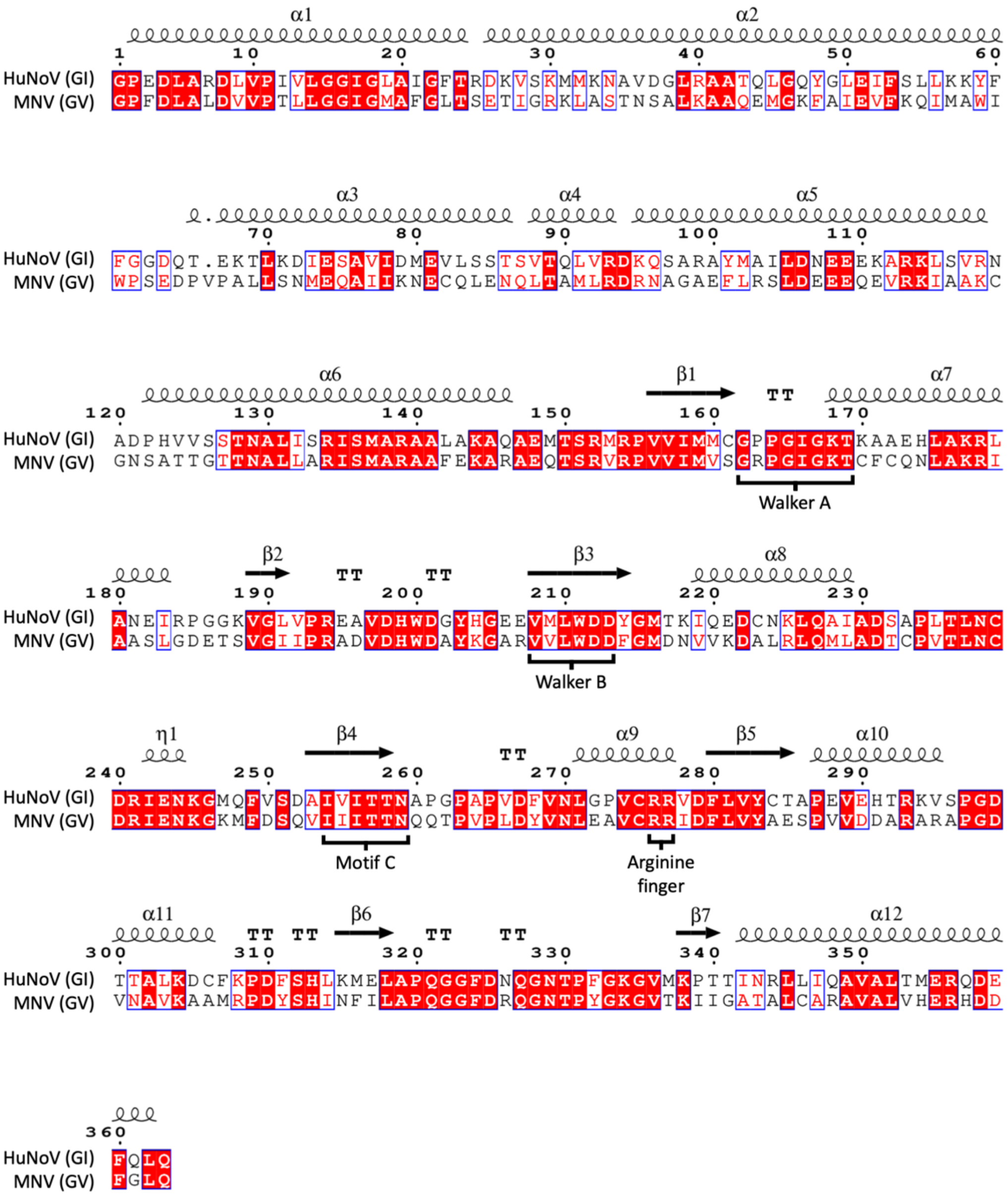

**Figure S9: Sequence alignment of HuNoV and MNV NS3 sequences.** Sequence alignment of HuNoV GI (NCBI Reference Sequence: NP_056820.1) and MNV GV (NCBI Reference Sequence: YP_724457.1) NS3 amino acid sequences. The sequence alignment was performed using Clustal Omega[9] and the image was generated by ESPript[10]. The secondary structure assignment, based on the HuNoV AlphaFold3 prediction, is shown. Signature SF3 helicase motifs are labeled.

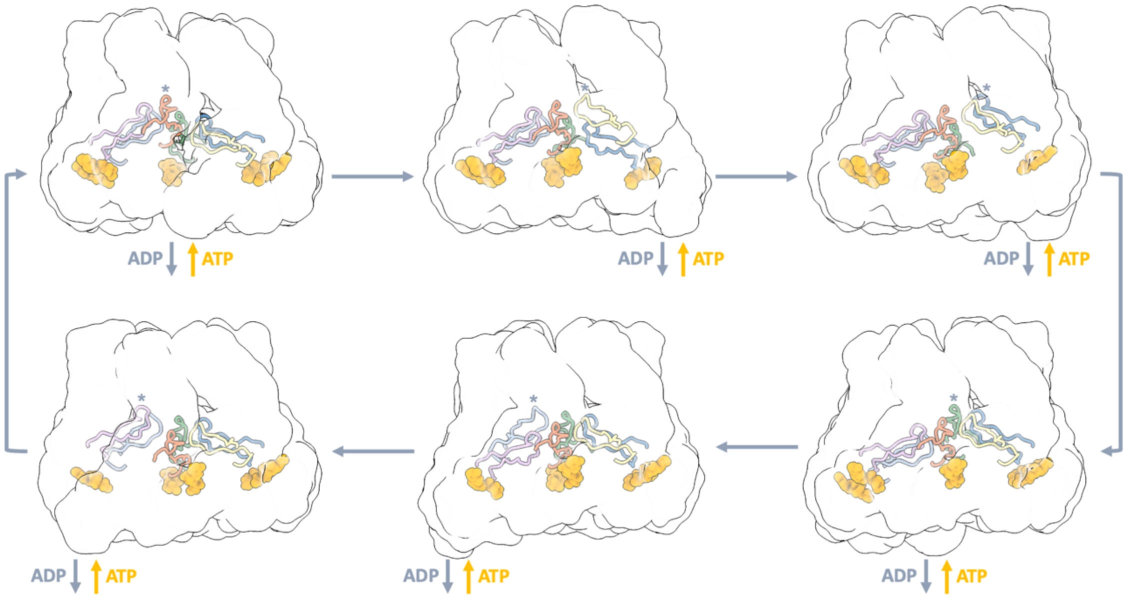

**Figure S10: Proposed model for NS3 ATP hydrolysis.** The six images depict different steps in the ATP hydrolysis cycle of NS3. The ATP binding site at the seam of the complex, where ADP is released and a new ATP molecule binds, is shown empty. Binding of an ATP to the nucleotide binding site of the subunit at position 6 (bottom of staircase) promotes its translocation to position 1 (top of the staircase). Consequently, each subunit moves to the next position in the staircase. For each step in the cycle, the uppermost pore loop 2 is indicated by an asterisk.

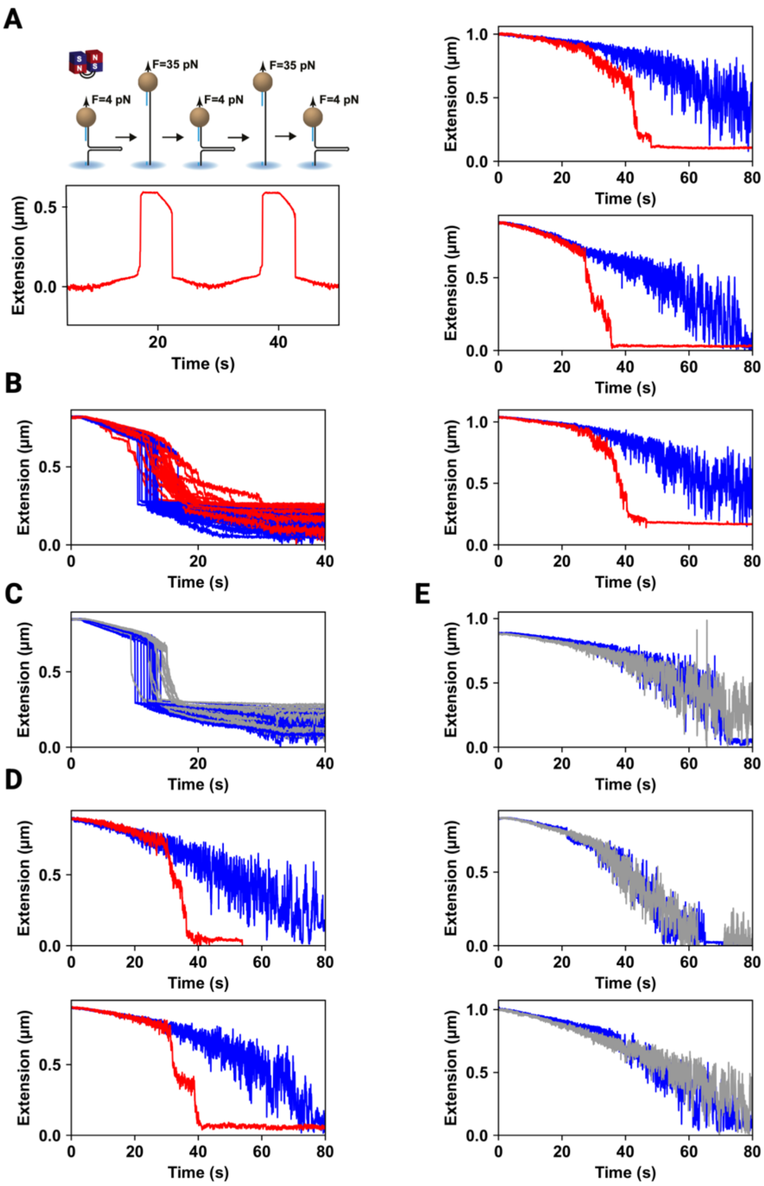

**Figure S11: (A) Hairpin characterization experiment preceding the experiments with NS3.** The force ramp shows the typical 1-step increase in extension of the hairpin upon reaching the critical force. (B, C) Hairpin refolding traces showing a pronounced delay in refolding in the presence of 1 µM hex-NS3 (red) compared to 1 µM MBP-hex (grey), overlaid on top of traces without the respective proteins (blue). After opening the hairpins by increasing the force to 35 pN, the force was subsequently lowered to 4 pN by moving the magnets away from the magnetic beads at a velocity of 0.1 mm/s (n = X for hex-NS3, n = Y for MBP-hex, n = Z for no protein). (D, E) Representative traces reporting dsRNA tether compaction in the absence (blue), or in the presence of 4.6 µM hex-NS3 (red) or 5 µM MBP-hex (grey). The magnets move slowly away from the flow cell surface at 0.1 mm/s. Experiments were performed in the absence of ATP. Individual bead measurements were n = 31 for hex-NS3 and n = 15 for MBP-hex.

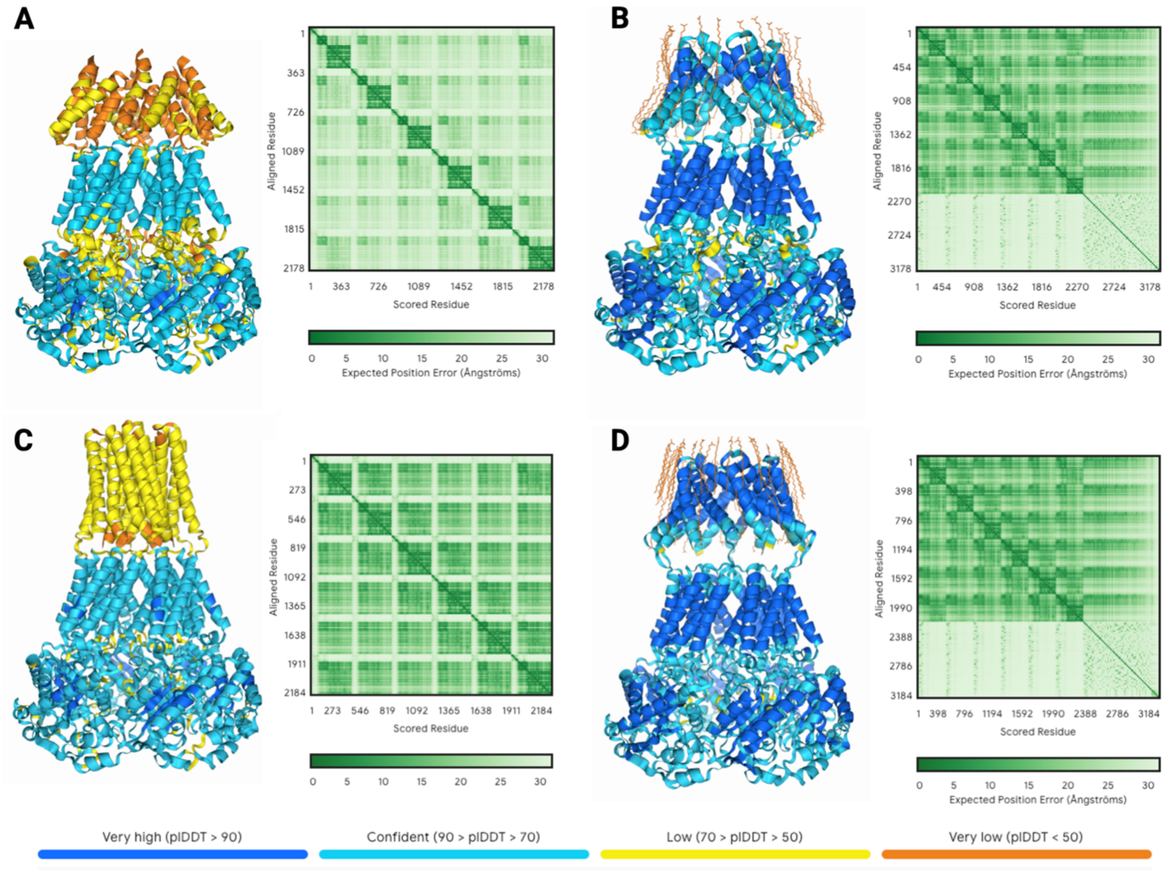

**Figure S12. AlphaFold3 predictions of GI and GV full-length NS3 hexameric complexes in the presence and absence of a membrane-mimicking environment.** (A) AlphaFold3[11] predicted structure of the GI NS3 hexamer. The protein backbone is shown in cartoon representation, colored by confidence score (predicted local distance difference test, pLDDT), with blue indicating high-confidence regions and red indicating low-confidence regions. The predicted alignment error (PAE) plot is included to assess the reliability of structural domains and interdomain orientations. (B) As in (A), but with the GI NS3 hexamer predicted in the presence of 50 molecules of oleic acid to mimic a membrane-like environment. (C) Predicted structure of the GV NS3 hexamer, displayed as in (A). (D) As in (C), but with the GV NS3 hexamer predicted in the presence of 50 molecules of oleic acid.

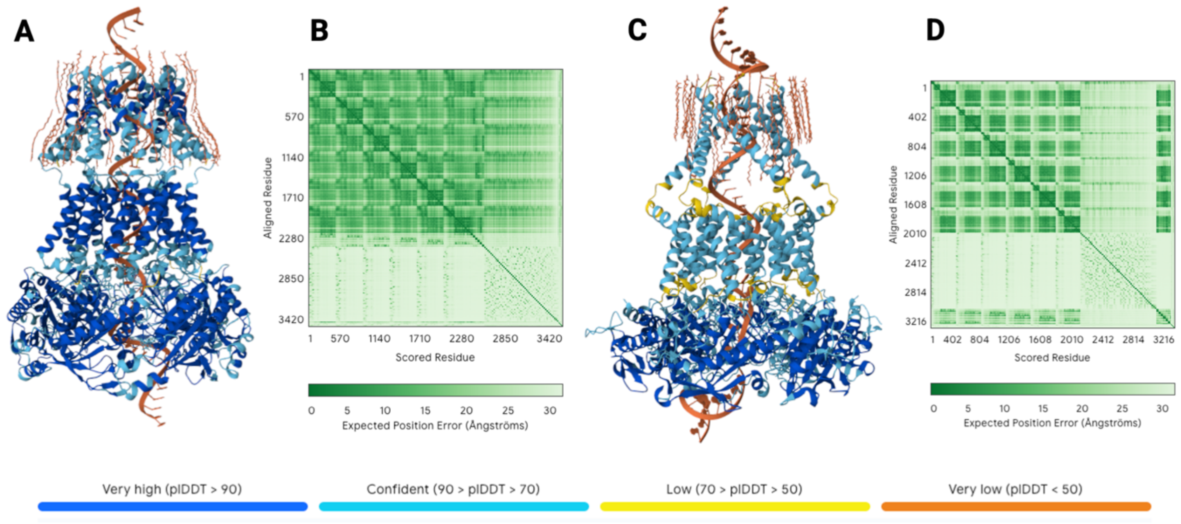

**Figure S13. AlphaFold3 predictions of the full-length GI NS3 and Coxsackievirus B3 2C hexameric complexes in the presence of a membrane-mimicking environment, RNA, and co-factors.** (A) AlphaFold3[11] predicted structure of the GI NS3 hexamer in complex with 50 molecules of oleic acid (to mimic a membrane environment), a 50-mer poly(A) RNA strand, 6 ATP molecules, and 6 magnesium ions. The protein backbone is shown in cartoon representation, colored by confidence score (predicted local distance difference test, pLDDT), with blue indicating high-confidence regions and red indicating low-confidence regions. The predicted alignment error (PAE) plot is included to evaluate the reliability of domain orientations and structural regions. (B) As in (A), but for the full-length Coxsackievirus B3 2C hexamer under identical conditions (50 oleic acid molecules, 50-mer poly(A) RNA, 6 ATP molecules, and 6 magnesium ions).

**Supplementary Table 1:** Data collection, image processing and refinement information

**Cryo-EM data collection, refinement and validation statistics**

| | HuNoV NS3 (EMDB-53547) (PDB 9R34) (EMPIAR-12758) |
|---|---|
| **Data collection and processing** | |
| Magnification | 130,000x |
| Voltage (kV) | 200 |
| Electron exposure (e–/Å$^2$) | 50.5 |
| Defocus range (μm) | 0.75-1.5 |
| Pixel size (Å) | 0.9 |
| Symmetry imposed | C1 |
| Initial particle images (no.) | 1,229,229 |
| Final particle images (no.) | 123,643 |
| Map resolution (Å) | 2.9 |
| FSC threshold | 0.143 |
| Map resolution range (Å) | 2-34 |
| **Refinement** | |
| Initial model used (PDB code) | AlphaFold2 |
| Model resolution (Å) | 2.8 |
| FSC threshold | 0.5 |
| Model resolution range (Å) | 2.6-2.8 |
| Map sharpening *B* factor (Å$^2$) | 64 |
| Model composition | |
| Non-hydrogen atoms | 14225 |
| Protein residues | 1800 |
| Ligands | 12 |
| *B* factors (Å$^2$) | |
| Protein | 41.8 |
| Ligand | 23.4 |
| R.m.s. deviations | |
| Bond lengths (Å) | 0.004 |
| Bond angles (°) | 0.584 |
| Validation | |
| MolProbity score | 1.59 |
| Clashscore | 4.9 |
| Poor rotamers (%) | 1.84 |
| Ramachandran plot | |
| Favored (%) | 97.32 |
| Allowed (%) | 2.68 |
| Disallowed (%) | 0 |

**Supplementary Video 1.**

Surface representation of the integrative model of the full-length HuNoV NS3 complex embedded in a membrane bilayer (left), and a central cross-section (right) highlighting a putative RNA translocation pathway. This animation illustrates the sequential ATP-driven conformational changes of the NS3 hexamer, consistent with a hand-over-hand translocation mechanism. Partially created with BioRender.com.
